# Feedstock-Efficient Conversion through Hydrogen and Formate-Driven Metabolism in *Escherichia coli*

**DOI:** 10.1101/2025.05.30.657096

**Authors:** Robert L. Bertrand, Justin Panich, Aidan E. Cowan, Jacob B. Roberts, Lesley J. Rodriguez, Juliana Artier, Emili Toppari, Edward E. K. Baidoo, Yan Chen, Christopher J. Petzold, Graham A. Hudson, Patrick M. Shih, Steven W. Singer, Jay D. Keasling

## Abstract

Product yields for biomanufacturing processes are often constrained by the tight coupling of cellular energy generation and carbon metabolism in sugar-based fermentation systems. To overcome this limitation, we engineered *Escherichia coli* to utilize hydrogen gas (H₂) and formate (HCOO⁻) as alternative sources of energy and reducing equivalents, thereby decoupling energy generation from carbon metabolism. This approach enabled precise suppression of decarboxylative oxidation during acetate growth, with 86.6 ± 1.6% of electrons from hydrogen gas (via soluble hydrogenase from *Cupriavidus necator* H16) and 98.4 ± 3.6% of electrons from formate (via formate dehydrogenase from *Pseudomonas* sp. 101) offsetting acetate oxidation. Hydrogen gas supplementation led to a titratable and stoichiometric reduction in CO₂ evolution in acetate-fed cultures. Metabolomic analysis suggests that this metabolic decoupling redirects carbon flux through the glyoxylate shunt, partially bypassing two decarboxylative steps in the TCA cycle. We demonstrated the utility of this strategy by applying it to mevalonate biosynthesis, where formate supplementation during glucose fermentation increased titers by 57.6% in our best-performing strain. Flux balance analysis further estimated that 99.0 ± 2.8% of electrons from formate were used to enhance mevalonate production. These findings highlight a broadly applicable strategy for enhancing biomanufacturing efficiency by leveraging external reducing power to optimize feedstock and energy use.

**Graphical Abstract:** 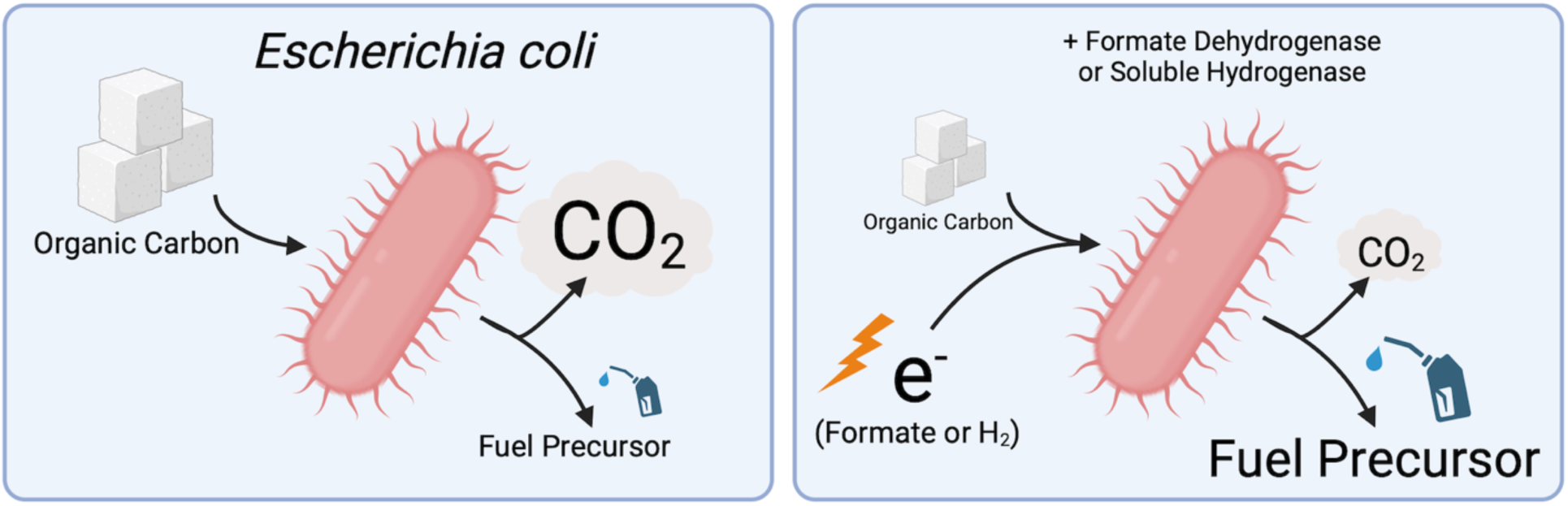

Created in BioRender. Panich, J. (2025)

https://BioRender.com/0tmtp94

## INTRODUCTION

Biomanufacturing—the microbial production of chemicals and materials—provides an alternative to conventional chemical synthesis, which often depends on high temperatures, extreme pressures, and toxic catalysts (Cowan et al., 2023; Zhang et al. 2017). Despite significant advances, biomanufacturing has had difficulty competing with the petroleum-based chemical industry due to higher production and feedstock costs (Keasling et al., 2021; Levi and Cullen, 2018). Bridging this economic gap necessitates novel strategies to optimize microbial energy and carbon utilization. In heterotrophic systems, cellular energy is primarily obtained from the oxidation and decarboxylation of carbon feedstocks (e.g., glucose). The feedstock, therefore, serves a dual role as both a substrate for biosynthesis and a source of reducing power for energy generation. These mutually exclusive roles present a seemingly intractable problem: every molecule of feedstock oxidized to generate ATP and NAD(P)H cannot also be transformed into a desired end product (Meadows et al., 2016; Ng et al., 2015; Tashiro et al., 2015). While numerous metabolic engineering approaches have been explored to enhance conversion and reduce carbon loss (Cui et al., 2016; Kang et al., 2016; Lin et al., 2018; Meadows et al., 2016; Ng et al., 2015; Tashiro et al., 2015), a fundamental challenge remains: how can we meet cellular energy demands without compromising product yield? Addressing this bottleneck is critical if industrial microbiology is to become economically competitive.

The emergence of electrolysis technology and the growing availability of inexpensive electricity allow for the exploration of biomanufacturing platforms that provide cells with external sources of reducing power. Feeding chemical electron donors to microbial hosts is not a new idea. However, the traditional approach involves identifying bioproducts with NAD(P)H-dependent biosynthetic pathways and then increasing their yield by improving NAD(P)H availability, either by supplying hosts with a sacrificial NAD(P)H donor or by rewiring host metabolism to reduce off-target NAD(P)H consumption (Liu et al., 2018; Wang et al., 2017). Essentially, NAD(P)H is viewed as limiting, and improved bioproduction is achieved through substrate-product kinetics or metabolic flux optimization (e.g., Balzer et al., 2013; Shen et al., 2011). In contrast, we observe that when energy is plentiful, cells downregulate feedstock oxidation via catabolic pathways (Lupacchini et al., 2021; Cotton et al., 2020; Sauer et al., 1999). Therefore, we hypothesize that providing an external source of reducing power can free feedstock from its traditional role as cellular fuel, increasing the amount of carbon that is available as substrate for bioconversion. Whether others see NAD(P)H as limiting, we invert this paradigm and instead view feedstock as limiting, then ask how it can be rescued from oxidative catabolism. This alternative framework is already used in microbiology to improve microbial growth efficiency (g_CDW_/g_SUBSTRATE_) (Babel, 2009). However, demonstrations of the direct causal link between external reducing power and enhanced product biosynthesis—by preventing feedstock oxidation—are lacking. Since all biological systems are compelled to oxidize feedstock, a key implication is its broad applicability: Electron donors should be able to enhance the production of any target molecule, and as will be shown, this even includes NAD(P)H-independent pathways.

Electron donation can be accomplished by equipping common industrial hosts with a mechanism for indirect electron transfer through electron carriers such as formate (HCOO^−^) or hydrogen gas (H_2_) (Claassens et al., 2018; Partipilo et al., 2023). Formate is readily oxidized by NAD^+^- or NADP^+^-specific formate dehydrogenase (FDH) to regenerate NADH or NADPH, and numerous structurally distinct FDH homologs are available to meet various performance criteria (catalytic efficiency, thermostability, proteolytic resistance, etc.). Formate is highly soluble, membrane-permeable, and can be electrochemically produced from CO_2_. However, it is toxic at high concentrations and requires pH buffering (Cotton et al., 2020; Yishai et al., 2016). NADH could also be regenerated by oxidizing H_2_ via hydrogenases. The sole byproduct of this reaction is a proton, which becomes covalently bound once NAD(P)H is re-oxidized (such as in respiration), thereby providing total atom economy and pH neutrality. Furthermore, H_2_ is non-toxic and energy-dense by unit mass. However, H_2_ is energy sparse by unit volume, poorly soluble in water, and explosive when mixed with oxygen (between 5-85% H_2_).

To investigate the effects of adding external reducing equivalents on metabolism, we have engineered *E. coli* to express the O_2_-tolerant soluble hydrogenase (HYD) from the chemolithoautotrophic microorganism *Cupriavidus necator* H16. In parallel experiments, we also used the FDH from *Pseudomonas sp. 101*. Both strains showed decreased feedstock consumption and increased product titer (mevalonate), indicating a broad potential for biomanufacturing.

## MATERIALS AND METHODS

### DNA Manipulations

Routine PCR was performed using Phusion Polymerase Hot Start II (*ThermoFisher*, Waltham, MA, USA) under the following conditions: 1 minute at 94°C; 25 cycles of 10 seconds at 94°C, 15 seconds at T_m_ - 4°C, and 20 seconds per kb at 72°C; followed by an additional 20 seconds per kb at 72°C. Amplicons were resolved by agarose gel electrophoresis, evaluated using gel imaging (*Bio-Techne*, Minneapolis, MN, USA), and purified with the QIAquick gel extraction kit (*Qiagen*, Hilden, Germany). Plasmid construction was carried out either by Gibson assembly with the NEBuilder HiFi DNA Assembly Master Mix *(New England Biolabs*, Ipswich, MA, USA) or through ligation using T4 DNA Ligase (*ThermoFisher*, Waltham, MA, USA). Candidate constructs were cloned using chemically competent “Stellar” *E. coli* (*Takaro Bio*, San Jose, CA, USA), and then authenticated either by nanopore sequencing (*Plasmidsaurus*, San Francisco, CA, USA) or by Sanger sequencing (*Azenta*, Burlington, MA, USA). Verified plasmids were subsequently transformed into DE3-competent BW25113 *E. coli* via electroporation. Plasmids (Table S1), primers (Table S2), bacterial strains (Table S3), and the plasmid assembly scheme (Figure S1) are available in the SI.

### Growth Media and Conditions

Hydrogenase prototyping experiments were conducted using Miller LB medium (*Millipore*, Burlington, MA, USA) and sterilized by autoclaving (hereafter “LB”). Acetate consumption assays utilized a rich media solution of MOPS, amino acids, and nucleotides (EZ Rich Defined Medium Kit: *Teknova*, Hollister, CA, USA), containing 55.5 mM sodium acetate, and was sterilized by 0.2-µm membrane filtration (hereafter “EZ Acetate”). Adding 5 g/L glucose instead of acetate produces “EZ Glucose.” Mevalonate production assays were carried out using M9 minimal medium containing 5 g/L glucose, 0.12 g/L MgSO_4_, 0.011 g/L CaCl_2_, 65 µg/L NiCl_2_, and a trace elements mixture containing (100x) 5 g/L EDTA, 0.83 g/L FeCl_3_-6H_2_O, 84 mg/L mM ZnCl_2_, 13 mg/L CuCl_2_-2H_2_O, 10 mg/L CoCl_2_-2H_2_O, 10 mg/L H_3_BO_3,_ and 1.6 mg/L MnCl_2_BO_3_-4H_2_O. This medium was sterilized by 0.2 µm membrane filtration (hereafter “M9 Glucose”). A bacterial washing step was performed using the same M9 minimal medium, but with glucose omitted (hereafter “M9 Wash”).

### Gas Cultivations

Glycerol stocks of BW25113 and CM15 *E. coli* were plated on LB agar with the appropriate antibiotics (*Teknova*, Hollister, CA, USA). Colonies were seeded into 10 mL of either LB, EZ Acetate, or M9 Glucose, and grown aerobically in culture tubes overnight at 37°C and 200 RPM for <18 h. Experimental cultures were established by inoculating 200 µL of seed cultures into 10 ml of the same medium (LB, EZ Acetate, or M9 Glucose) in glass serum bottles. Cultures were grown mid-logarithmically by aerobic cultivation at 37°C and 150 RPM for 4 h. The cultures were then transferred to a 30°C warm room. Hydrogenase expression was induced using 0.25 mM IPTG, whereas formate dehydrogenase expression was constitutively expressed. Bottles were closed with butyl stoppers and crimped with aluminum to create an airtight seal. From the approximately 112 mL headspace, 20 mL of atmospheric air was removed using a syringe fitted with a one-way stop valve. The cultures expressing hydrogenase were then supplied with either 20 mL of H_2_ gas (800 μmol) or 20 mL of N_2_ gas. The H_2_ addition step is accurate to ± 5 μmol H_2_ (Table S4 in *SI*). The cultures expressing formate dehydrogenase were given either 40 mM sodium formate or a water blank and otherwise equivalently treated by withdrawing 20 mL of air, followed by adding 20 mL of N_2_ gas. Cultures were grown for 18 h at 150 RPM and 30°C. To enhance H_2_ uptake during mevalonate assays when *E. coli* was cultivated in glucose minimal medium, a variant procedure was performed as follows: Seed cultures were inoculated (200 µL) into 10 mL of EZ Glucose in glass serum vials and grown aerobically at 37°C and 150 RPM for 4 h. Cultures were then transferred to a 30°C warm room, induced with 0.25 mM IPTG, and allowed to grow aerobically for four more hours. Cells were centrifuged and resuspended in M9 Wash. Cells were again centrifuged and then resuspended in M9 glucose (5 g/L) containing 0.25 mM IPTG. Serum vials were stoppered and crimped, and 20 mL of air was replaced with 20 mL of H_2_ or N_2_. Cultures were incubated for 18 h at 150 RPM and 30°C. Culture density was evaluated using optical density (OD_600_) and/or dry cell weight (g/L). The cellular dry weight (CDW) was measured by concentrating an aliquot of bacterial suspension within pre-weighed microcentrifuge tubes, followed by drying for 48 hours. The consensus OD600/CDW ratio was 3.07 ± 0.17, in agreement with prior studies (Sauer et al. 1999). The CDW was measured in most experiments, and actual CDW readings are shown in figures wherever possible. In cases where only OD_600_ data were collected, these readings are converted into CDW using the consensus ratio, as indicated in figure captions. Diagrams of key experimental schemes are provided (Figures S2 to S4 in the *SI*). Statistical replicate cultures of BW-HYD grown in EZ-Acetate revealed statistical variations of 1% to 4% with respect to OD_600_, cellular dry weight (CDW), CO_2_ production, and broth acetate concentration (Table S5 in *SI*).

### Gas Analysis

The GoDirect Gas Pressure Sensor (*Vernier*, Beaverton, OR, USA) was used to monitor the internal gas pressure of serum bottles. Broth aliquots for downstream analyses (e.g., HPLC) were collected either using a needle and syringe or by opening serum vials with needle-nose pliers and removing aliquots by pipetting. Gas analysis of dissolved CO_2_ was enabled by acidifying media aliquots with a few drops of concentrated HCl. Gas analysis was conducted using a Shimadzu GC-2014 gas chromatograph, withdrawing gas aliquots from butyl-stoppered serum vials with syringes fitted with one-way stop valves. Samples flowed isothermally through a 2 mm x 8 m Shimadzu (Kyoto, Japan) column at 60°C using argon as the carrier gas. H_2_, O_2_, and N_2_ were quantified after 8.5 minutes of flow time through a TCD module at 120°C, with CO_2_ quantified after an additional 4.0 minutes of flow time through the methanizer at 380°C and FID module at 200°C. The gas profile was quantified by interpolating peak integrations to standard plots, using inert N_2_ as an internal standard. Gasses are quantifiable by GC with an error of ± 0.21% (Table S6 in SI).

### *In Vitro* Hydrogenase Assays

Cells were inoculated from an overnight pre-culture in LB with appropriate antibiotics and induced for hydrogenase expression at 30°C during the mid-log phase. Cells were cultivated for 24 or 48 h, cold centrifuged, and lysed by sonication. Total protein content was evaluated using the Bradford assay. Hydrogenase activity from lysates was assessed with an NADH absorption assay at 340 nm, as previously described (Lupacchini et al., 2021). This work was performed in an anaerobic growth chamber containing 4% H_2_ and 10% CO_2_, and assays were analyzed using a 340PC SpectraMax spectrophotometer (*Molecular Devices,* San Jose, CA, USA).

### Extracellular Metabolite Quantification

The profiles of extracellular metabolites were evaluated using HPLC. Samples were prepared by isolating the broth supernatant and adding 10% methanol to create an internal loading standard. The samples were filtered through 0.45-µm modified nylon centrifugal filters (*VWR*, Radnor, PA, USA), and the filtrates were stored at −80°C until needed. The samples were analyzed isothermally on an Aminex HPX-87H column (*BioRad*, Hercules, CA, USA) set to 65°C, with a refractive index detector set to 50°C (*Agilent*), and the sample loading tray maintained at 8°C (*Agilent*, Santa Clara, CA, USA). The mobile phase consisted of 0.005 M H_2_SO_4_ and flowed at a rate of 0.6 ml/min. Data acquisition and analysis were performed using ChemStation software (*Agilent*, Santa Clara, CA, USA). The broth concentrations of glucose, formate, ethanol, acetate, glycerol, lactate, succinate, and mevalonate were quantified by interpolating peak integrations against authenticated standards (*Sigma-Aldrich*, St. Louis, MO, USA). Acetate is quantifiable by HPLC with an error margin of ± 2.2 μmol (Table S7 in *SI*).

### Additional Materials and Methods

Methods for LC-MS, protein identification and quantification, HPLC-MS, computational flux balance analysis, and deductive metabolic estimations are found in the *SI*.

## RESULTS

### Prototyping the *C. necator* soluble hydrogenase (HYD) and *Pseudomonas sp.* 101 formate dehydrogenase (FDH)

NAD^+^-dependent hydrogenases reversibly reduce the nicotinamide cofactor with electrons from H_2_ (Lubitz et al., 2014). While most hydrogenases are sensitive to O_2_ and are inactivated upon contact with air (Lu and Koo, 2019), a key exception is the soluble and O_2_-tolerant hydrogenase from *C. necator* H16 (formerly *Ralstonia eutropha* H16) (Pandelia et al., 2013), an organism that has drawn attention for its ability to use energy gained from H_2_ to aerobically fix CO_2_ and transform it into bioplastics and biofuels (Panich et al., 2021; 2024). The soluble hydrogenase is a heterotetramer (HoxFUYH), and its heterometallic NiFe catalytic core and iron-sulfur clusters require nine maturation proteins (HoxWI and HypABCDEFX) to assemble.

The expression of 13 soluble proteins in a heterologous host presents a considerable challenge. However, it has already been functionally expressed in *E. coli,* providing a catalog of genetic components (Ghosh et al., 2013; Lonsdale et al., 2015; Fan et al., 2022; Teramoto et al., 2022; Lamont and Sargent, 2017; Schiffels et al., 2013). The existing plasmids and expression systems were non-optimal due to missing genes, non-codon-optimized genes, and multiple plasmids. Using the genetic components described in Lamont and Sargent (2017), and kindly donated by them to us, we consolidated all 13 genes into a single plasmid containing the medium copy p15A origin of replication and kanamycin resistance, organized into two operons (T5 and *tatA* promoters) (Figure S1 in *SI* for assembly scheme and Figure S5 in *SI* for plasmid map). Transforming this plasmid into *E. coli* BW25113 resulted in strain BW-HYD.

Cultivating BW-HYD with H_2_ and air in sealed bottles demonstrated hydrogenase’s functional activity, directly by gas chromatography (Figure 1A) and evidenced indirectly by pressure monitoring (Figure 1B) in LB media. We complemented these experiments with metabolomics analyses to investigate any effects that H_2_ may have on BW-HYD metabolism. Cultures treated with H_2_ displayed increased cellular concentrations of reduced metabolites of the TCA cycle (succinate, fumarate, malate) compared to N_2_-treated controls (Figure S6). We hypothesize that cells respond to H_2_ uptake by engaging the glyoxylate shunt, a carbon-conserving bypass of the TCA cycle that reduces carbon loss as CO_2_ by minimizing decarboxylative NADH generation (Dolan and Welch, 2018). This hypothesis is supported by biosynthetic studies utilizing the glyoxylate shunt to enhance the production of succinate, fumarate, malate, and other closely related intermediates (Yang et al., 2022).

**Figure 1:**
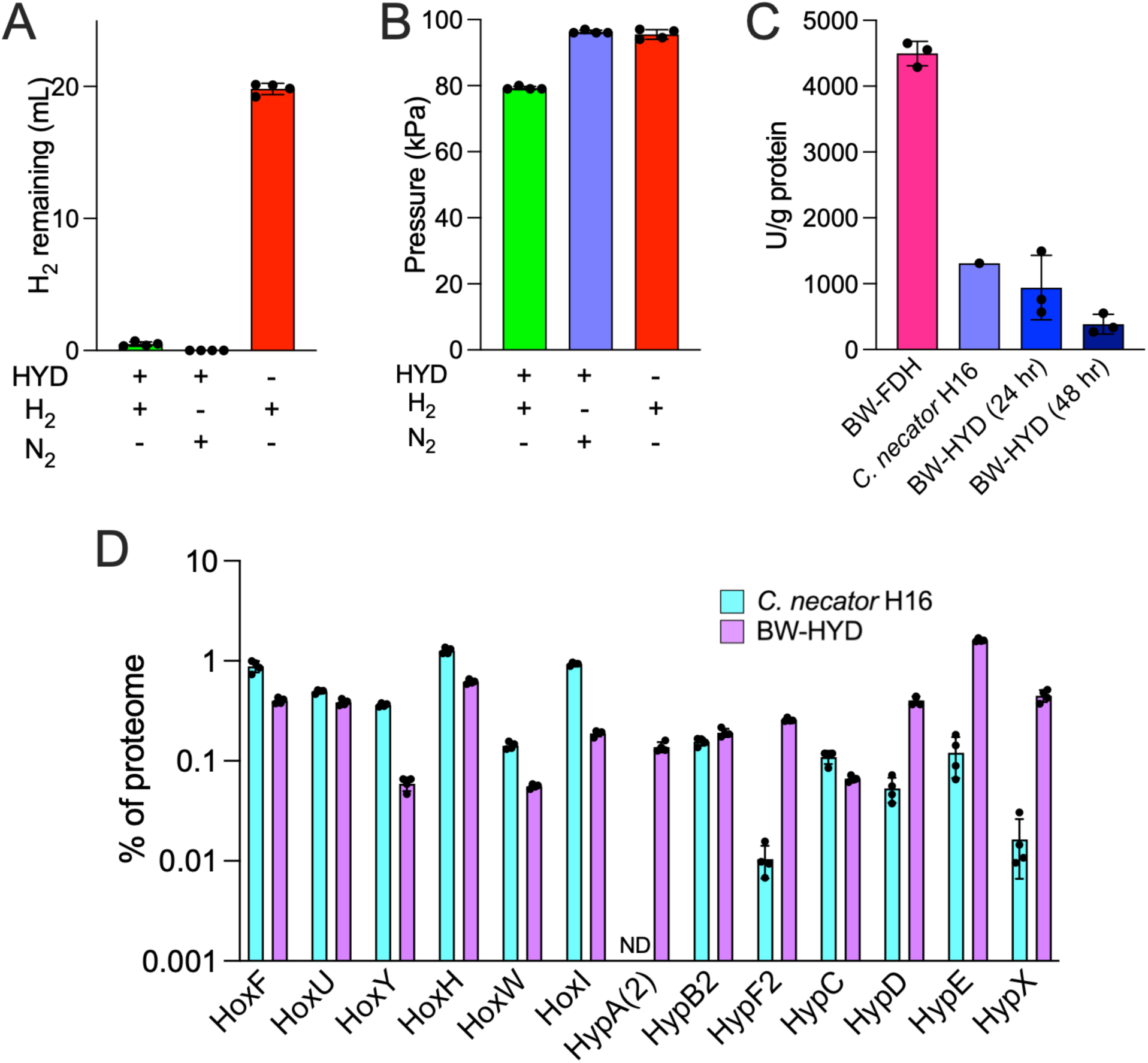
Characterizing the heterologous soluble hydrogenase from *C. necator* H16 in *E. coli*. **(A)** H_2_ consumption (expressed as ml H_2_ remaining in headspace) by *E. coli* BW25113 expressing *C. necator* hydrogenase (BW-HYD), when initially provided with 20 ml H_2_ mixed with air, compared to the same strain provided with 20 ml N_2_ mixed with air, and untransformed BW25113 fed with 20 ml H_2_ mixed with air, after an overnight incubation in sealed serum bottles containing approximately 112 ml headspace. **(B)** Final internal pressure of enclosed serum vials of these same cultures. **(C)** Specific *in vitro* activity measurements of *C. necator* hydrogenase within crude lysates of BW-HYD after 24h or 48h of cultivation (n=3), compared to this same hydrogenase when expressed within the native producer (*C. necator* H16) grown in 62% H_2_ / 10% CO_2_ / 10% O_2_, balanced with N_2_ (n=1). The specific activity of the *Pseudomonas sp.* FDH on formate was used as a reference control when grown for 24 h (n=3) (Gleizer et al., 2019). An enzymatic unit (U) is 1 µmol substrate per minute. **(D)** Proteomics analysis to quantify the protein abundance of the 13 *C. necator* hydrogenase heterotetramer and maturation proteins when expressed in *E. coli*, in comparison to their natural expression levels in *C. necator* (n=4). HypA is denoted HypA(2) because HypA1 and HypA2 sequences are identical; ND = Not detected. HypA(2) was not detected in *C. necator* samples, which may reflect very low abundance below detection limits or peptide misassignment during analysis. See Figure S5 in the *SI* for the plasmid map.

We then performed biochemical characterization experiments to compare the specific activity of our construct in *E. coli with* lysates from autotrophically grown *C. necator* H16. Cells were cultured for the designated times, lysed, and analyzed for NADH production in the presence of saturating NAD^+^ (Lupacchini et al., 2021). The hydrogenase activity observed from BW-HYD was similar to the activity measured in *C. necator* (942 U/g CDW and 1310 U/g CDW, respectively), aligning with previous reports of soluble hydrogenase activity in *C. necator* H16 (Schlesier and Friedrich, 1981) (Figure 1C). Compared to lysates collected 24h after inoculation, hydrogenase activity in BW-HYD decreased after 48h of cultivation. We hypothesize that this decline is due to gradual enzyme denaturation or instability.

In addition, we performed quantitative proteomics on the 13 heterologous proteins expressed in *E. coli* and compared these results with those from autotrophically grown *C. necator*. We observed that the expression of HoxY in *E. coli* is substantially lower than that of HoxFUH, indicating an unbalanced expression of the heterotetramer (Figure 1D). Moreover, BW-HYD produced many maturation proteins at higher levels than those in *C. necator*, suggesting that maturation proteins are expressed in excess. Cumulatively, these experiments demonstrate that a prototype plasmid encoding the *C. necator* hydrogenase and its maturation machinery was successfully constructed and functionally expressed in *E. coli*, but that expression stoichiometry should be considered during future optimization efforts, as more cellular energy appears to be expended assembling proteins than is minimally required.

To broaden the scope of this study, we employed a formate dehydrogenase (FDH) as an alternative mechanism for assimilating external electrons. We selected the FDH from *Pseudomonas sp.* 101, which has been extensively characterized in *E. coli* (Gleizer et al., 2019; Tishkov et al., 1993; Wenk et al., 2020). Transforming a plasmid encoding this FDH into *E. coli* BW25113 yielded strain BW-FDH. In cell lysates, this FDH exhibited over a fourfold increase in specific activity for NADH formation compared to HYD (Figure 1C). The superior performance of FDH could relate to its simpler expression and maturation requirements compared to hydrogenases.

### Electron donors prevent the decarboxylation of an organic feedstock

The oxidation of H_2_ and HCOO^−^ results in the regeneration of NADH. Either donor should satisfy cellular energy requirements because NADH can drive oxidative phosphorylation or transfer its hydride equivalent to regenerate NADPH through the native transhydrogenase PntAB. We tested both donors’ ability to inhibit the oxidative decarboxylation of feedstock by cultivating BW-HYD on acetate. Acetate was chosen because NADH production from this feedstock depends solely on the TCA cycle, which simplifies experimental interpretation (Wenk et al., 2020).

BW-HYD was cultivated in EZ Rich (Takara) with 55.5 mM acetate in lieu of glucose (see experimental scheme in Figure S2A of *SI*). This is a defined medium composed of amino acids, nucleotides, and inorganic nutrients, of which acetate is the predominant carbon source at about 50% by mass. Mid-logarithmic phase cultures were provided with increasing amounts of H_2_ (0 to 800 µmol). After an overnight (16h) incubation in butyl-stoppered serum vials, the effects of H_2_ uptake on biomass formation, acetate retention, and CO_2_ production were evaluated. Results show that all cultures consumed most (>80%) of the H_2_ supplied (Figure S7 in *SI*) and that H_2_ consumption did not alter how much biomass formed, thereby normalizing this variable (Figure 2A). A linear molar relationship between H_2_ uptake and acetate retention was observed (3.925 ± 0.062; Figure S8B in *SI*), demonstrating that BW-HYD consumed less acetate to produce the same amount of biomass. This improved efficiency coincided with a decrease in CO_2_, indicating that less feedstock was oxidized (Figure 2B). According to stoichiometry, in order to prevent the formation of two moles of CO_2_, it is necessary to also decrease the oxidation of one mole of acetate. This stoichiometry was observed (2.035 ± 0.036; Figure S8A in *SI*), providing direct evidence that the oxidation of hydrogen gas was replacing the oxidation of acetate as a means of generating cellular energy.

**Figure 2:**
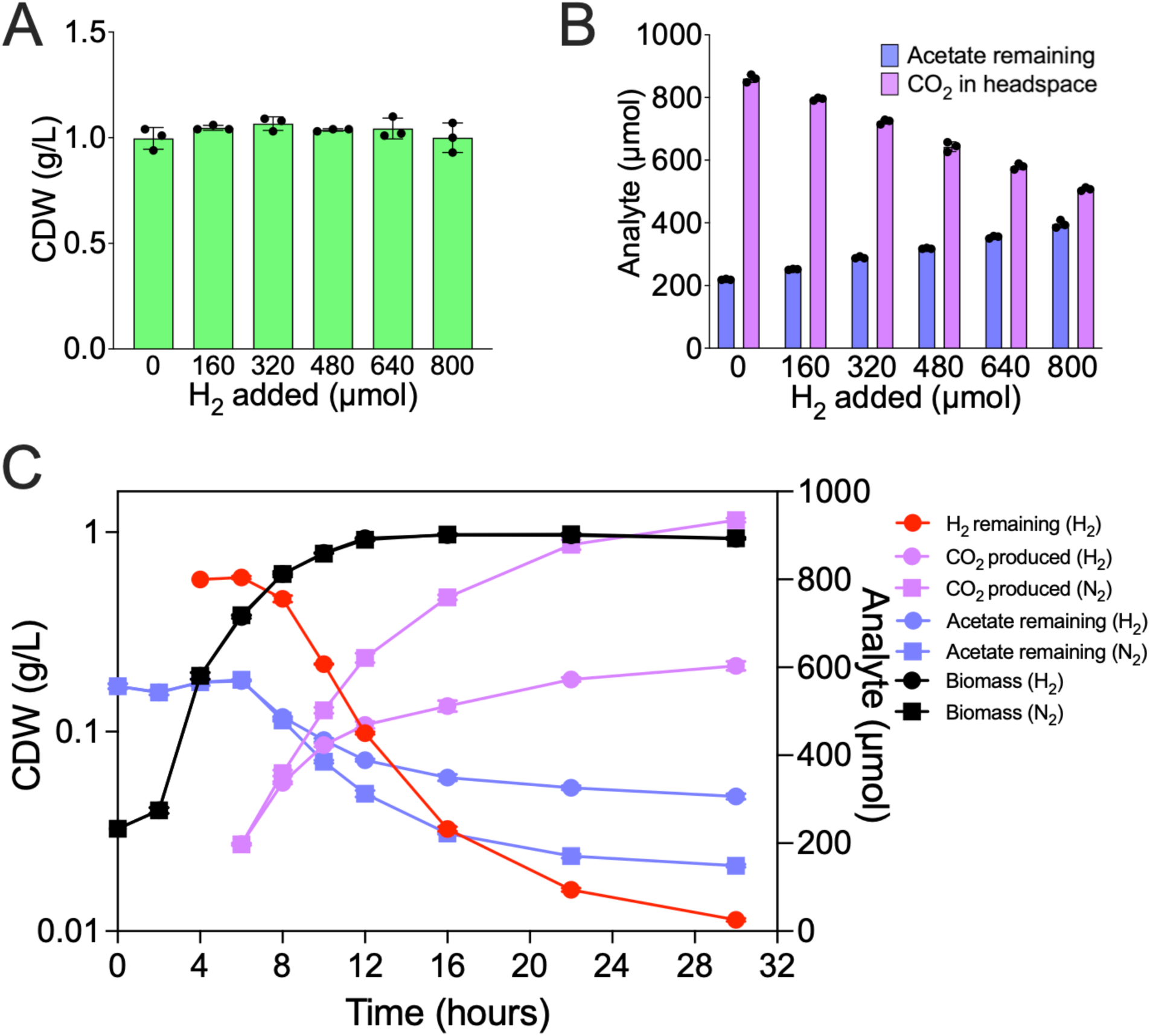
Titrating the metabolic effects of H_2_ uptake on acetate consumption and CO_2_ production in BW-HYD. **(A)** Biomass formation (dry cell weight) after an overnight incubation (n=3). Hydrogen gas uptake data are shown in Figure S7 in the *SI*. **(B)** Acetate remaining in the culture media (µmol) and carbon dioxide produced (µmol) (n=3). Plots of H_2_ expended versus acetate retained, and of CO_2_ avoided versus acetate retained, are available in Figure S8 in the *SI*. **(C)** A time course experiment evaluating the effects of H_2_ uptake on acetate retention and CO_2_ production by BW-HYD, compared to the same strain subjected to an inert N_2_ treatment. Either 20 ml of H_2_ (800 µmol) or 20 ml of N_2_ was added to the headspace of each culture at time = 4 hours. Biomass accumulation is shown in black, measurements of H_2_ are shown in red, acetate measurements in solution are shown in blue, and CO_2_ measurements are shown in purple. Circles represent cultures incubated in air and fed with H_2_, while squares represent cultures incubated in air and treated with an equivalent volume of inert N_2_. The culture optical density (OD_600_) was recorded and converted into CDW using the consensus ratio (see *Gas Cultivations*). The experiment was performed in technical triplicate using a common seed culture inoculum (n=3). Some error bars are occluded by symbols.

Feedstock preservation was also shown through a time-course assay. H_2_ uptake, biomass formation, and acetate consumption were measured over 30 hours, during which BW-HYD cultures were grown in 55.5 mM acetate, supplied with either 20% H_2_ in air or 20% N_2_ in air, and harvested at specific times (see experimental scheme in Figure S3 in *SI*). Changes in H_2_ uptake, CO_2_ production, and acetate consumption appeared four hours after IPTG induction, indicating that *E. coli took* about four hours to translate and activate the hydrogenase (Figure 2C). The peak H_2_ consumption rate was 8.4 to 9.4 mmol H_2_ / g_CDW_ / h and was seen 8-12h post-inoculation. Acetate concentration dropped at 8 h, coinciding with the start of H_2_ uptake and differences in CO_2_ production, suggesting a causal link. No differences in lag phase length or relative growth rates between H_2_- and N_2_-treated cultures were observed, showing H_2_ enhances feedstock use efficiency but does not change growth kinetics. Cellular replication rates can be roughly estimated by comparing culture densities at different times. The average doubling time before acetate consumption was 1.05h (up to 6h post-inoculation), the ten hours that followed presented with an average doubling time of 5.75h (6h to 16h post-inoculation). This suggests biphasic growth typical of diauxic growth. This explains why biomass increased without acetate consumption in the first six hours, as cells likely consumed amino acids, nucleotides, and other components of the rich media first, followed by acetate as a secondary substrate. Biphasic growth is also supported by O_2_ data (Figure S9A in *SI*), as peak O_2_ consumption was observed 6h post-inoculation at 12.9 mmol O_2_ / g_CDW_ / h, thereafter dropping at 10h by one-third to 8.8 mmol O_2_ / g_CDW_ / h despite continued cell growth, and fell to 3.3 mmol O_2_ / g_CDW_ / h once stationary phase has started (16h).

While H_2_ is appealing because of its low cost and energy density, formate is a non-explosive energy carrier that can similarly power microbial metabolism and may have broader utility. To this end, we performed an equivalent end-point titration using sodium formate (see experimental scheme in Figure S2B in *SI*). Formate can be oxidized by FDH into NADH and CO_2_ using the BW-FDH strain, which was cultivated similarly to BW-HYD. In triplicate cultures, increasing amounts of formate (0 to 400 µmol) were fed to BW-FDH during the mid-logarithmic phase. Cultivation in butyl-stoppered serum vials enabled CO_2_ measurements. Following the experiment, formate was undetectable, suggesting that 100% of the formate was consumed under all tested conditions (Figure S10A in *SI*). As observed with BW-HYD grown with H_2_, biomass formation was unaffected by formate addition, normalizing cellular energy requirements among triplicates (Figure 3A). A titratable molar relationship (2.865 ± 0.103; Figure S8D in *SI*) emerged between formate addition and acetate retention. We distinguish between “biogenic CO_2_” arising from the TCA-mediated oxidation of carbohydrate feedstocks such as acetate and “abiogenic CO_2_” arising from the FDH-mediated oxidation of formate. The oxidation of formate to regenerate NADH produces abiogenic CO_2_ as a byproduct, which complicates our assessment of the relationship between biogenic CO_2_ emissions and acetate oxidation (Figure S10B in *SI*). Therefore, to determine moles of biogenic CO_2_, we subtracted moles of formate oxidized from the total amount of CO_2_ observed in the headspace (Figure 3B). This calculation is appropriate because *E. coli* BW25113 is incapable of autotrophic growth on CO_2_: If abiogenic CO_2_ decreases due to fixation (e.g., by PEP carboxylase), this must coincide with an equal rise in biogenic CO_2_ production (or else biomass accumulation on CO_2_ becomes possible). A stoichiometric relationship (1.933 ± 0.110; Figure S8C in *SI*) was revealed between biogenic CO_2_ abatement and acetate preservation, demonstrating that this alternative electron donor is also capable of suppressing the oxidation of a feedstock.

**Figure 3:**
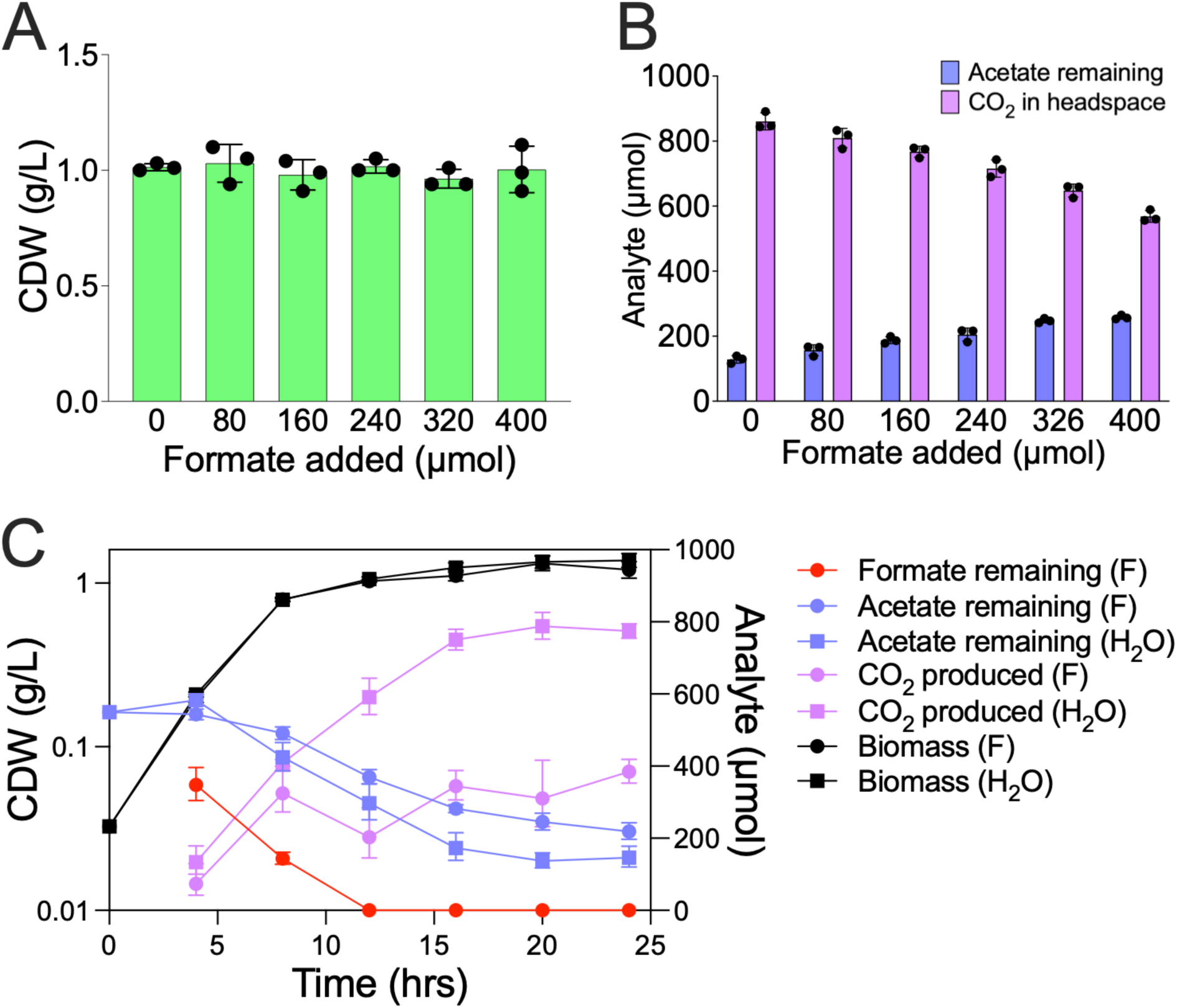
Titrating the metabolic effects of formate uptake on acetate retention and CO_2_ production in BW-FDH. **(A)** Biomass formation at the end of an overnight incubation (dry cell weight) (n=3). These data show that formate uptake did not influence the amount of biomass formed, thereby normalizing biomass as a variable when examining the effects of formate uptake on acetate consumption and CO_2_ production. Formate uptake data are presented in Figure S10 in the *SI*. **(B)** Acetate remaining in the culture media and biogenic CO_2_ produced (n=3). Biogenic CO_2_ is calculated by subtracting the moles of formate oxidized from total CO_2_ (µmol). Total CO_2_ production can be found in Figure S10B in the *SI*. **(C)** A time course experiment evaluating the effects of formate uptake on acetate retention and CO_2_ production by BW-FDH (F), compared to the same strain subjected to an inert water treatment (H_2_O). Either 400 µmol or an equivalent volume of ddH_2_O was added to each culture at time = 4 hours. Biomass accumulation is shown in black, measurements of formate are shown in red, acetate measurements in solution are shown in blue, and biogenic CO_2_ measurements are shown in purple (CO_2_ from formate oxidation was subtracted from measured values of CO_2_). Circles represent cultures incubated in air and fed with formate (F), while squares represent cultures incubated in air and treated with an equivalent volume of water (H_2_O). The culture optical density (OD_600_) was recorded and converted into CDW using the consensus ratio (see *Gas Cultivations*). The experiment was performed in technical quadruplicates using a common seed culture inoculum (n=4). Some error bars are occluded by symbols.

To further assess the kinetics of formate utilization, we performed a time-course assay comparing BW-FDH cultures supplemented with 400 µmol formate or water blanks as a control (Figure 3C and Figure S9B in *SI*). Formate-fed cultures rapidly and completely oxidized the added donor, consistent with endpoint titration results (Figure S10A). Acetate consumption slowed immediately after formate addition, while biomass accumulation remained unchanged, demonstrating that electrons from formate effectively substituted for acetate oxidation in meeting cellular energy demands. After correcting for abiogenic CO_2_ derived from formate oxidation, we observed that biogenic CO_2_ production was consistently lower relative to controls. Notably, divergence between treated and untreated cultures emerged within hours of supplementation, indicating a rapid metabolic shift upon the availability of exogenous reducing power. Compared to hydrogen-fed BW-HYD (Figure 2C), where uptake was delayed by the time required for enzyme maturation, FDH-enabled formate oxidation occurred immediately, underscoring the advantage of formate as a soluble, membrane-permeable donor that can be assimilated without delay.

### Electron donors increase mevalonate yield with tunable efficiency

Our results so far show that electron donors can replace an organic feedstock as the source of cellular energy needed to produce biomass. We then explored whether providing H_2_ and HCOO^−^ to *E. coli* could enhance the production of a specific bioproduct. Feedstock is typically a limiting factor for heterotrophic biomanufacturing because all biological systems must oxidize carbon to generate cellular energy. Therefore, augmenting NADH accumulation should improve bioproduction.

We investigated this possibility using mevalonate (MEV), the key metabolic intermediate in isoprenoid biosynthesis and a precursor in the production of high-value commodity chemicals. It was chosen because its biochemical pathway is well-understood and it can be easily measured with HPLC using commercially verified standards. While most MEV pathways are NADPH-specific, we instead utilized the NADH-specific HMG-CoA reductase from *Delftia acidovorans* (Kang et al., 2019). When paired with NADH-specific HYD and NADH-specific FDH, this enzyme allows direct coupling of chemical electron donors to MEV production.

To investigate, we inserted plasmids encoding hydrogenase or formate dehydrogenase, along with genes for NADH-specific mevalonate biosynthesis (Table S1 in *SI*), resulting in strains BW-HYD-MEV and BW-FDH-MEV (Table S3 in *SI*). We faced two procedural challenges. First, initial experiments using the same acetate-based rich medium as in the previous section showed a baseline mevalonate titer of only 30 mg/L (Figure S12 in *SI*), a low value that hampers our ability to clearly distinguish differences in titer between treatments and controls. The low mevalonate titer was likely due to acetyl-CoA being necessary for MEV biosynthesis, and it is well-known that acetyl-CoA is limited in cells grown on acetate because of competing demands from central carbon metabolism and enzymatic bottlenecks such as acetyl-CoA carboxylase (Jeung et al., 2023). In contrast, glucose is widely used in industry to produce commodity chemicals because its metabolism generates excess acetyl-CoA and NADH. To overcome this obstacle, we used glucose as the feedstock, observing that growing BW-HYD-MEV and BW-FDH-MEV in glucose (5 g/L) M9 minimal media consistently produced baseline MEV titers between approximately 0.5 and 1.0 g/L (Figure 4). The second challenge was that H_2_ consumption by BW-HYD was negligible when applying the same cultivation protocol from the previous section, which involved inducing hydrogenase and introducing H_2_ at 4h post-inoculation (Figure S2A in *SI*). We anticipated that the protocol would need adjustment because H_2_ uptake by exogenous hydrogenases varies due to genetic background and cultivation conditions (Teramoto et al., 2022; Maeda et al., 2007). The results in Figure 2C offered two clues for protocol development: (1) hydrogenases require 4 hours to mature after IPTG induction, and (2) strong hydrogenase expression occurs in rich media. To address this, we tested a modified protocol where BW-HYD was first inoculated into rich medium and grown for 4 hours to mid-log phase. Hydrogenase expression was induced with IPTG, and cells were grown for another 4 hours. The cells were then washed, transferred to glucose minimal medium, and cultivated overnight in defined headspaces of 20% H_2_ in air or 20% N_2_ in air (control condition) (Figure S4B in *SI*). This approach proved successful, as BW-HYD was now consuming approximately 15 of the 20 mL of H_2_ provided—a promising result for future application on BW-HYD-MEV (Figure S13 in *SI*). Unlike hydrogenase, no uptake issues were observed for formate when growing *E. coli* in glucose minimal medium, so a simpler 4h post-inoculation protocol was used for BW-FDH-MEV (Figure S4B in *SI*).

**Figure 4:**
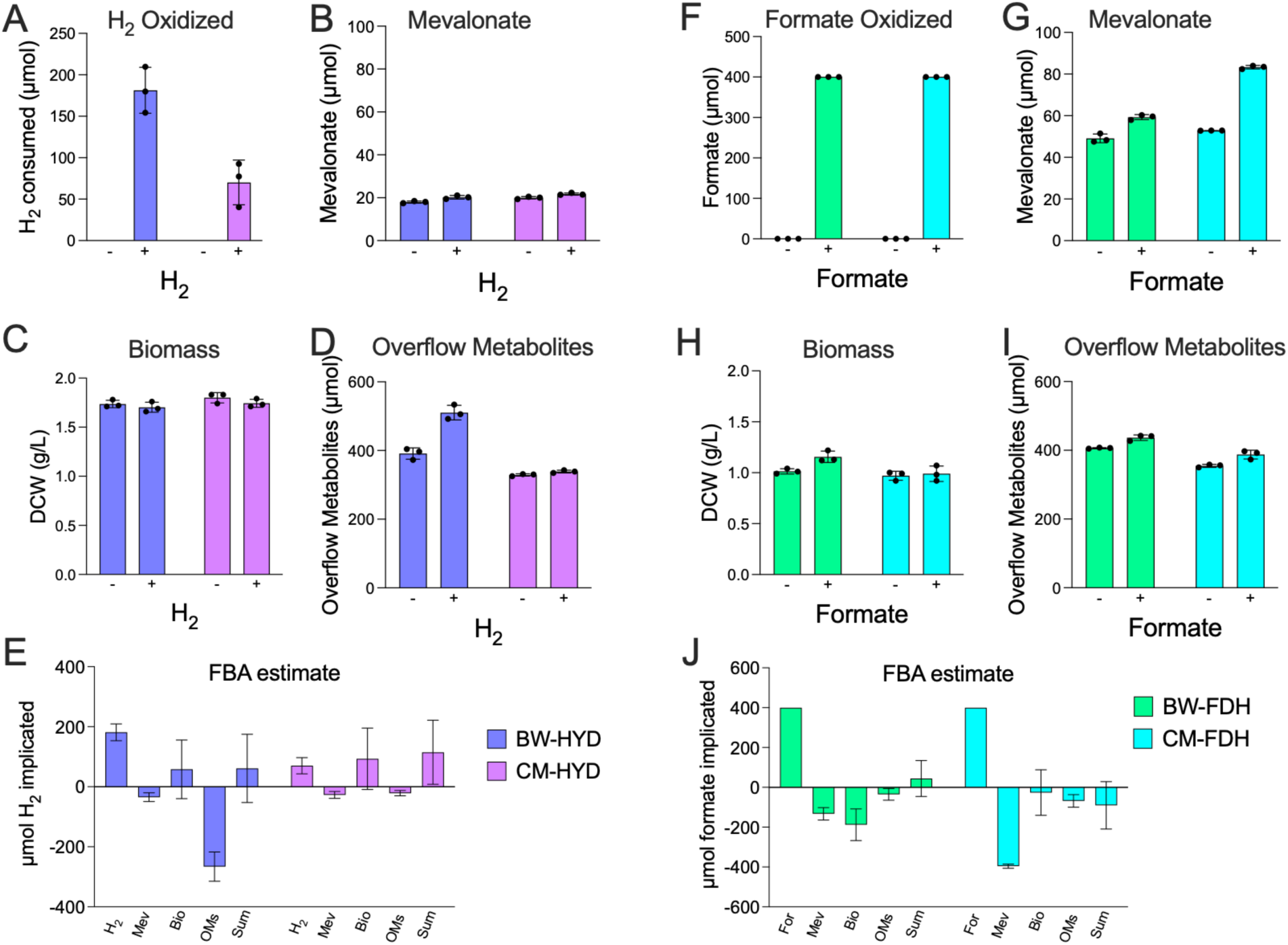
Effects of hydrogen gas (A-E) and formate (F-J) on the biochemical profiles of BW-HYD-MEV (blue bars), CM-HYD-MEV (purple bars), BW-FDH-MEV (green bars), and CM-FDH-MEV (cyan bars) in glucose M9 minimal media. **(A, F)** Amounts of formate and hydrogen gas consumed (µmol). **(B, G)** Mevalonate produced (µmol). **(C, H)** Dry cell weight (g/L). **(D, I)** Profile of mixed acid production (µmol), summed by the moles of carbon atoms contained within formate, ethanol, acetate, glycerol, lactate, and succinate; micromoles of each mixed acid are detailed in Table S8 in *SI*. **(E, J)** Estimates from computational flux balance analysis (FBA) of the hydrogen gas or formate required to produce each biochemical difference observed between electron-treated and blank-treated cultures, where negative values indicate the amount of hydrogen gas or formate that must be theoretically oxidized to produce that change, and positive values suggest either the donation of electrons or its equivalence in surplus cellular energy. Only acetate, ethanol, and formate (in hydrogenase-laden cultures) were analyzed for fermentation impact because these three molecules comprise >95% of the mixed acid profile. The sum of positive and negative contributors is expected to fall within the margin of zero micromoles, as discussed in *the flux balance analysis to evaluate the efficiency of hydrogen gas and formate usage*. Abbreviations: Mev = Mevalonate; Bio = Biomass; OMs = Overflow Metabolites.

The results of these parallel experiments are shown in Figure 4. BW-HYD-MEV utilized 181 μmol of the 800 μmol H_2_ in headspace (Figure 4A). Despite this limited uptake, there was a modest increase in MEV titer from 18.0 ± 0.5 to 20.2 ± 0.8 μmol (0.266 to 0.299 g/L) (Figure 4B). These data suggest that 82.3 moles of H_2_ were oxidized for each mole of surplus MEV produced. Although minimal differences in biomass formation were noted (Figure 4C), H_2_ prompted the production of C_1_ (formate) and C_2_ (ethanol and acetate) mixed acid products (Figure 4D; Table S8 in *SI*). Compared to BW-HYD-MEV, all 400 μmol of formate supplied to BW-FDH-MEV were fully consumed, as shown by the absence of detectable formate in the culture supernatant after overnight cultivation (Figure 4F). Formate oxidation elevated mevalonate production from a baseline titer of 49.1 ± 2.1 to 59.3 ± 1.2 μmol (0.73 to 0.88 g/L) (Figure 4G), providing a ratio of 39.2 moles of formate oxidized per mole of surplus MEV generated. Sample readings revealed that the addition of formate also enhanced the production of biomass (Figure 4H) and overflow metabolites (Figure 4I). These experiments demonstrate that both electron donors can boost the production of a commodity chemical, but much of the extra reducing power was redirected to generate surplus biomass and overflow metabolites.

We hypothesized that blocking pathways responsible for mixed acid production would redirect carbon flux toward mevalonate synthesis, thereby increasing its production. We tested this hypothesis using *E. coli* CM15 (MG1655 *ΔaraBAD ΔfadRE ΔldhA ΔackApta ΔadhE ΔpoxB ΔfrdABCD*) (Mehrer et al., 2018), creating equivalent strains CM-HYD-MEV and CM-FDH-MEV by introducing the appropriate plasmids (Table S3 in *SI*). Hydrogen gas uptake by CM-HYD-MEV was modest at only 70 μmol H_2_ (Figure 4A). We suspect that this lower H_2_ uptake results from the strain lacking fermentative pathways that would typically serve as redox sinks. Despite this, the baseline MEV titer increased from 20.0 ± 0.5 to 21.7 ± 0.5 μmol (0.296 to 0.322 g/L) (Figure 4B). This results an H_2_/MEV molar ratio of 41.2, a twofold improvement compared to the ratio of 82.3 observed with BW-HYD-MEV. The consumption of 400 μmol of formate by CM-FDH-MEV (Figure 4F) significantly raised the baseline titer from 52.9 ± 0.1 to 83.3 ± 0.8 μmol (0.78 to 1.23 g/L) (Figure 4G). This represents a threefold increase in the formate/mevalonate molar ratio, from 39.2 (BW-FDH-MEV) to only 13.2. Notably, using CM15 minimized differences in biomass and mixed acid formation between treatment and control conditions (Figures 4C, 4D, 4H, 4I), indicating that a larger share of electrons from formate and H_2_ was directed toward mevalonate biosynthesis. This rerouting aligns with a host unable to dispose of excess reducing power through standard fermentation. Interestingly, excess biomass accumulation was also limited in CM15-based strains. We hypothesize that the absence of flexible redox balancing pathways prevents these strains from using surplus electrons for growth, despite increased energy availability, although the exact mechanism remains unclear. These findings demonstrate that electron donors can promote the production of a commodity chemical and that donor utilization efficiency can be improved through host strain engineering.

To support the generalizability of increasing bioproduction yields using chemical electron donors, we also tested flaviolin, an NAD(P)H-independent polyketide produced by RppA (Yang et al., 2018; Incha et al., 2020). Supplying H_2_ to BW-HYD when expressing RppA increased flaviolin production by about 30% compared to N_2_-treated controls (Figure S11), indicating that electron donors can also enhance bioproducts that do not directly consume NADH. Owing to technical limitations, including the lack of a commercially available standard, we discontinued flaviolin studies in favor of the quantitative mevalonate system. This preliminary experiment nonetheless suggests that chemical electron donors may have broad applicability.

We have observed that cultures treated with H_2_ respond by increasing the production of succinate, fumarate, and malate (Figure S6 in *SI*). The rise in these three metabolites suggests activation of glyoxylate shunt metabolism (Dolan and Welch, 2018; Yang et al., 2022). We hypothesize that engagement of the non-oxidative, carbon-retaining glyoxylate shunt pathway could explain how feedstock retention, CO_2_ avoidance, and bioproduct formation are possible (Figures 2, 3, 4). The glyoxylate shunt might explain how a microbial host can increase feedstock usage efficiency while maintaining redox balance (Figure 5). This hypothesis is supported by observations that shunt activation increases acetate accumulation and reduces CO_2_ production (Waegeman et al., 2011; Noronha et al., 2000).

**Figure 5.**
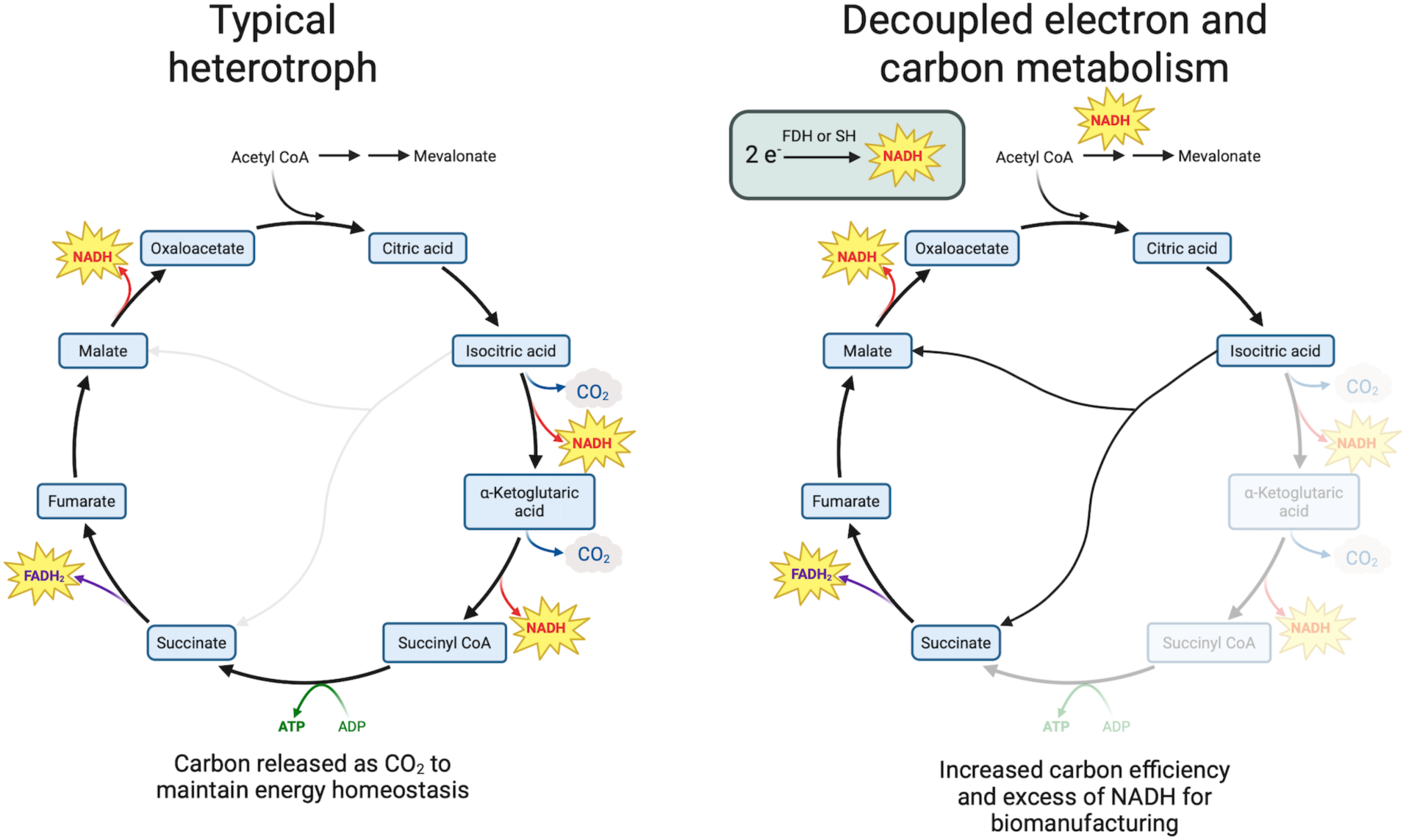
The proposed mechanism for carbon conservation in cells engineered with FDH or HYD is to increase carbon flux through the glyoxylate shunt. Created in BioRender. Panich, J. (2025): https://BioRender.com/l26rpn5

### Flux balance analysis to evaluate the efficiency of hydrogen gas and formate usage

We used flux balance analysis (FBA) to evaluate the efficiency of hydrogenase and FDH in producing reducing equivalents that would otherwise be derived from oxidizing acetate. FBA is a systems biology technique that utilizes stoichiometric relationships within metabolic networks to compute feasible flux distributions under various constraints without the need for kinetic data. We used FBA to predict the marginal yield envelope of *E. coli,* defined as the range of potential changes in product yield in response to alterations in a specific substrate when supplemented with either hydrogen gas or formate. Typically, FBA operates on an optimization principle, such as maximizing the molar yield of a product in response to a specific substrate and is based on the assumption that evolution has optimized metabolism to achieve optimal yields. This assumption is valid in model organisms like *E. coli* when cultivated on acetate and glucose, as the observed fluxes and growth rates align with FBA predictions (Varma and Palsson, 1994). However, this premise may not hold when using synthetic metabolic pathways, as these pathways are not subject to evolutionary optimization pressures. A limitation of FBA is its presumption that organisms always strive for maximal yield, whereas experiments with *S. cerevisiae* and *C. necator indicate* that organisms may also prioritize pathways that sacrifice yield for faster growth rates or metabolic flexibility (Schuster et al., 2008; Jahn et al., 2021). To validate the computational FBA findings, and as described in the *SI*, we also utilized a heuristic and deductive reasoning approach to estimate molar ratios, where the key step involves equating all energy carriers (ATP, NAD(P)H, FADH_2_) to proton motive force. As shown in Table 1, the computational FBA and deductive estimates closely align.

**Table 1:** Compared to untreated *E. coli*, how many moles of hydrogen gas or formate need to be oxidized to….

| Metabolic effect* | Hydrogen gas |  | Formic acid |  |
| --- | --- | --- | --- | --- |
|  | FBA | Estimate | FBA | Estimate |
| ...prevent the complete oxidation of one mole of acetate? | 3.40 | 3.25 | 2.82 | 2.89 |
| ...enable the formation of one additional gram of dry biomass from glucose? | 0.157 | 0.123 | 0.131 | 0.109 |
| ...enable the production of one additional mole of mevalonate from glucose via the mevalonate pathway? | 15.60 | 14.75 | 13.00 | 13.11 |
| ...enable the fermentation of one additional mole of acetate from glucose? | 3.60 | 3.75 | 3.00 | 3.33 |
| ...enable the fermentation of one additional mole of ethanol from glucose? | 6.20 | 6.25 | 5.17 | 5.56 |
| ...enable the fermentation of one additional mole of formic acid from glucose? | 2.00 | 1.125 | N/A<sup>a</sup> | 1.00 |

<sup>a</sup> Modeling the effects of formic acid addition upon formic acid fermentation resulted in a negative value. \*"FBA" is computational flux balance analysis, as described in the main text. "Estimate" uses deductive reasoning to estimate bioenergy requirements based on well-established biochemical facts, as described in SI.

The efficiency of H_2_ or formate in preventing the consumption of acetate (Figures 2 and 3), and of enhancing mevalonate production (Figure 4), can be determined by the following equation:

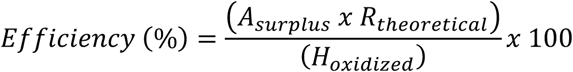

Where:

*A*_surplus_ = Moles of surplus acetate (Fig. 2B, 3B) or surplus mevalonate (Fig. 4B, 4G)
*R*_theoretical_ = Theoretical moles of H_2_ or formate required per mole of surplus A (Table 1)
*H*_oxidized_ = Moles of H_2_ (Fig. S7, 4A) or formate (Fig. S10, 4F) oxidized in culture

We have demonstrated a titratable and stoichiometric relationship between H_2_ consumption, acetate retention, and CO_2_ avoidance (Figure 2; Figure S8 in *SI*). The number of moles of H_2_ that are experimentally required to avoid the oxidation of one mole of acetate can be determined by plotting moles of H_2_ consumed against moles of acetate retained. The resultant molar ratio is 3.925 (± 0.062) (Figure S8B). In contrast, FBA indicates that only 3.40 moles of H_2_ are needed (Table 1), suggesting that 86.6% (± 1.8) of the electrons supplied by H_2_ were used to replace acetate oxidation as a source of cellular energy. We speculate that the remaining 13.4% of the reducing power was used in off-target reactions not captured by the analysis. Compared to H_2_, fewer moles of formate (2.82) are theoretically required to achieve this effect because formate crosses the cell membrane as formic acid, with its proton contributing to the electrochemical potential (Table 1). Similar to H_2,_ we also performed a formate titration by incrementally increasing formate concentrations and quantifying acetate and CO_2_ responses (Figure 3; Figure S8 in *SI*). Plotting the formate consumption and acetate retention data revealed that 2.865 (± 0.102) moles of formate are required to prevent the oxidative decarboxylation of one mole of acetate (Figure S8D), suggesting that 98.4% (± 3.6) of the electrons supplied by formate successfully replaced acetate for cellular energy generation.

The efficiency of H_2_ or formate in enhancing the formation of mevalonate can be determined in four steps: (1) assess the moles of H_2_ or formate oxidized within cultures (Figures 4A, 4F); (2) measure the moles of surplus mevalonate formed within H_2_- or formate-treated cultures (Figures 4B, 4G); (3) multiply the surplus moles of mevalonate by the number of moles of H_2_ or formate that FBA determines is theoretically required to produce this surplus (Table 1); and (4) divide this amount by the total moles of H_2_ or formate oxidized within cultures and express the result as a percentage (maximum of 100%). Both H_2_ and HCOO^−^ increased mevalonate biosynthesis (Figures 4B and 4G). Mevalonate is produced by condensing three moles of acetyl-CoA, followed by NADH-dependent reduction. To boost mevalonate formation, electron donors must correspondingly prevent the decarboxylation of three moles of acetyl-CoA and meet this NADH requirement. Analysis of BW-HYD-MEV revealed that of the 181 (± 28) μmol H_2_ oxidized by this strain (Figure 4A), only 19.1% (± 8.1) was utilized to enhance mevalonate titer (Figure 4E). Interestingly, H_2_-fed cells had a slightly lower biomass quantity than control cells (Figure 4C): Compared to control cells, the energy not used for this missing biomass is equated to the consumption of an additional 58.2 (± 97.7) μmol H_2_ by H_2_-fed cells (Figure 4E). This shift in cellular energy usage explains how the surplus of overflow metabolites in H_2_-fed cells can require an energy input equivalent to 146.8% (± 26.9) of all H_2_ oxidized. The total amount of H_2_ theoretically required to facilitate these biochemical changes is thus 120 (± 113) μmol H_2_, which is within the error range of the 181 μmol H_2_ experimentally observed as used (See “SUM” in Figure 4E). As compared to H_2_, the FBA findings of BW-FDH-MEV were statistically robust: Of the 400 μmol formate oxidized, 33.2% (± 7.7) was employed to enhance mevalonate production, 46.9% (± 19.8) was utilized to increase biomass formation, and 8.8% (± 7.4%) was used to boost the production of mixed acid products (Figure 4J). Total biochemical changes relative to water-blanked cultures required the theoretical oxidation of 355 µmol (± 90 µmol) of formate, which aligns with the error of the 400 µmol of formate consumed (See “SUM” in Figure 4J).

Changing the base strain from BW25113 to CM15 resulted in improved mevalonate production efficiency in both CM-HYD-MEV and CM-FDH-MEV. In CM-HYD-MEV, H_2_-fed cells now devoted 37.8% (± 16.2) of the energy supplied by 70 μmol of oxidized H_2_ toward mevalonate formation. However, after considering biomass and mixed acid changes, we were unable to balance total energy requirements between treatment and control conditions (see “SUM” in Figure 4E), a result likely arising from low statistical resolution caused by low H_2_ uptake. In contrast to H_2_, performance in CM-FDH-MEV was exceptional: virtually all energy provided to CM-FDH-MEV through the oxidation of formate was utilized to enhance mevalonate titer (99.0 ± 2.8%) (see “SUM” in Figure 4J). In conclusion, these data demonstrate that all (within error) of the energy supplied to an industrial host by formate can be harnessed to boost the production of a commodity chemical. However, additional strain engineering will be necessary to improve both the consumption of H_2_ and the efficiency of this supplemental energy use.

## DISCUSSION

This study demonstrates a promising strategy to overcome a fundamental limitation in biomanufacturing by decoupling cellular energy generation from carbon metabolism through the use of external electron donors. We successfully engineered *E. coli* strains capable of utilizing hydrogen gas and formate as alternative energy sources, achieving remarkable efficiencies: 86.6 ± 1.8% of electrons from H₂ and 98.4 ± 3.6% of electrons from formate were effectively used to replace acetate oxidation for cellular energy needs. Most notably, we demonstrated that up to 99.0 ± 2.8% of formate-derived energy can be redirected toward enhanced bioproduct formation when coupled with appropriate metabolic engineering strategies. Our proposed mechanism of upregulated flux through the glyoxylate shunt provides a mechanistic framework for understanding how external reducing power enables simultaneous CO₂ reduction and enhanced substrate utilization. These findings collectively support the feasibility of “feedstock-efficient bioconversion,” where organic substrates serve exclusively as building blocks rather than dual-purpose energy sources and raw materials.

### Engineering Hydrogenase Expression and Activity

While our system achieved functional hydrogenase activity, several optimization opportunities emerged from detailed characterization. Why did cell lysates of BW-HYD present with comparable hydrogenase activity to *C. necator* lysates despite imbalanced heterotetramer expression (Figure 1C)? Comparative proteomics revealed that HoxY in BW-HYD was produced at about 1/10 the rate as HoxFUH, meaning that most of the HoxFUH produced was likely inert. We observe that many maturation proteins in *E. coli* were also produced in excess (∼10x) to native expression levels in *C. necator* (Figure 1C). It’s possible that maturase overexpression may have compensated for HoxY underexpression. This hypothesis is informed by recent efforts to clone the *C. necator* hydrogenase into the cyanobacterium *Synechocystis sp.*: Whereas low activity was observed and efforts to raise this activity by increasing heterotetramer expression were ineffective, activity rose sharply when maturases were robustly expressed (Lupacchini et al., 2025). Recently, Siebert and coworkers (2025) reported efforts to optimize *C. necator* hydrogenase activity in *E. coli*, of which their work relied upon the same set of genetic components that we used to build the single plasmid hydrogenase cassette (see Lamont and Sargent, 2017). Similar to us, they also observed low HoxY production, which they hypothesized was caused by low levels of maturase HoxW. They observed that increasing HoxW expression increased NAD⁺ reduction activity by 40% and improved subunit stoichiometry. These studies exemplify the importance of maturase engineering upon hydrogenase activity and provide a validated troubleshooting guide for heterotetramer balancing.

Notably, it is not necessarily true that all nine *C. necator* maturases must be expressed in *E. coli*: A hydrogenase cassette encoding only five maturation proteins was nonetheless functional in *E. coli*, presumably because endogenous maturases were able to functionalize the *C. necator* hydrogenase (Ghosh et al., 2013). This suggests the possibility of creating a minimal cassette encoding only those maturases that are strictly necessary for functional expression, which could be designed through a knockout trial and error process. Finally, whereas our plasmid used a medium copy origin of replication (P15A), nothing precludes designing low copy vectors or even single copy chromosomally integrated strains.

From a biomanufacturing perspective, plasmid replication, gene transcription, and translation of exogenous proteins all impose fitness costs. The application of relative proteomics empowered us with biologically actionable insights related to heterotetramer balancing. Nonetheless, future application of absolute quantification proteomics would reveal greater information, such as an estimation of the fraction of the total proteome allocated to hydrogenase machinery. However inefficient hydrogenase expression is at present, we may nonetheless deduce that the energy gained by H₂ must exceed the metabolic burden imposed by this enzymatic engine, or else acetate retention given identical biomass (Figure 2) and surplus mevalonate biosynthesis given identical feedstock (Figure 4) would not be possible.

We have estimated by flux balance analysis that 86.6 ± 1.8% of H₂ that was consumed by BW-HYD was used to prevent the oxidation of acetate (Figure 2). This suggests that at least some portion of the remaining 13.4% was expended to maintain the 13-member hydrogenase machinery, though the exact amount of H₂ remains unclear. In contrast, equivalent experiments applying formate upon BW-FDH revealed that 98.4 ± 3.6% was used to offset acetate oxidation. This difference in efficiency could be reasonable if we consider that formate dehydrogenase is a single protein compared to the complex 13-protein hydrogenase system. FDH appears to be more promising than hydrogenase in this activity in our experiments. While FDH is theoretically reversible, it is thermodynamically favored to operate in the formate-oxidizing direction as the Δ*E°’* of the reaction is roughly +110 mV (compared to H_2_/H^+^→NAD^+^/NADH *ΔE°’* = +94 mV). This exogonic property may play an outsized role in throughput, as we must consider that H_2_ mass transfer would limit the hydrogenase enzyme in the forward direction whereas CO_2_ mass transfer would instead limit FDH in the reverse reaction.

### Metabolic Regulation and Growth Dynamics

We quantified the proportion of total cellular energy demand that is satisfied by the consumption of H_2_. Data from Figure 2C revealed that peak hydrogen gas draw rate occurred at the 8-10h assay interval (9.4 mmol H₂ / g CDW / h) followed by the 10-12h assay interval (8.4 mmol H₂ / g CDW / h). If we approximate the dry mass of an *E. coli* bacterium to 200 femtograms (Loferer-Krößbacher et al., 1998; Heldal et al., 1985), assume an e⁻/H⁺ yield ratio of 4 (Sharma et al., 2012; Borisov et al., 2011), and an H⁺/ATP expenditure ratio of 4 (Steigmiller et al., 2008), then peak energy input is around 600,000 ATP / cell / second. In contrast, ATP reporter experiments have demonstrated that exponentially growing *E. coli* require 6,400,000 ATP / cell / second (Deng et al., 2021), implying that the hydrogenase is only supplying about 10% of all energy needed by *E. coli*. However, that study was conducted aerobically using glucose-rich and minimal media, whereas the *E. coli* of Figure 2C were instead grown on acetate in stoppered serum vials with diminishing oxygen partial pressure. Although doubling times of 20 to 30 minutes typify glucose-rich media (Lin & Jacobs-Wagner, 2022), we observed a doubling time of about 5h when peak H_2_ consumption occurred (8 to 12h post-inoculation). Therefore, the true ATP requirement is likely to be substantially lower than what is required during glucose cultivation.

The 16h to 30h interval of Figure 2C is informative because no further accumulation of biomass is observed (onset of stationary phase), but consumption of H_2_ and acetate persisted (ongoing cellular maintenance costs), thereby providing a steady-state condition. In this latter 14h stationary period, about 206 μmol of H_2_ were consumed, as was 43 μmol of acetate. The oxidation of one mole of acetate is equivalent in cellular energy to the oxidation of 3.4 moles of H_2_ (Table 1). We can therefore estimate that oxidizing 43 μmol of acetate provided a total volume of energy equal to 292 μmol of ATP over 14h. Similarly, the expected energy yield of oxidizing 206 μmol of H₂ is 412 μmol of ATP. The combined ATP contribution is 704 μmol, and when averaged over 14h, equals 177,000 ATP / cell / s. This addition is consistent with the known 200,000 ATP / cell / s consumption rate of stationary phase *E. coli* (Deng et al., 2021), and suggests that H_2_ is satisfying 58% of cellular energy demand, and acetate oxidation the remaining 42%. Alternatively, we may evaluate the ratio of acetate consumption in H_2_-fed cells (43 μmol) to the control cultures (76 μmol). From this ratio, we may estimate that H_2_ satisfies 43% of total demand. It would be reasonable to conclude that the true answer is somewhere around half, and we caution that this is merely an estimation.

Cultivation of BW-HYD in rich defined acetate media resulted in onset of stationary phase at around 16h (Figure 2C and SXA in *SI*). This occurred despite the availability of O_2_, H_2_, and acetate, ruling out anaerobic metabolism and organic feedstock exhaustion as plausible explanations. Growth arrest in the presence of carbon sources is nonetheless possible when non-carbon metabolites such as nitrogen, phosphate, sulfur, and magnesium are limiting (Chubukov & Sauer, 2014). Under such conditions, catabolic programs persist and cells continue to regenerate NAD(P)H by oxidizing carbon to CO_2_ (Rühl et al. 2012). This hypothesis would be consistent with our observations of ongoing acetate and H₂ oxidation as well as continued CO_2_ generation during the stationary phase (Figure 2C). Importantly, both chemical electron donors allowed *E. coli* to produce an identical amount of biomass while consuming less feedstock (Figures 2 and 3), demonstrating that electron donors decouple energy generation from feedstock oxidation and can increase biomass production efficiency without increasing biomass production volume.

### Carbon Source-Dependent Hydrogen Uptake and Mechanistic Insights

We reported difficulty achieving high H₂ consumption in BW-HYD (Figure S13) and BW-HYD-MEV (Figure 4) when grown in glucose-minimal media. In contrast, H_2_ uptake by BW-HYD was robust in acetate-rich media (Figure 2; Figure S7). In addition, no obstacles were reported when others have used this same enzyme in *E. coli* to ferment H_2_ using glucose as an electron source (Ghosh et al., 2013; Schiffels et al., 2013; Lamont & Sargent, 2017). We provide two possible explanations.

First, at pH 7.0, the thermodynamic potential of the 2H⁺/H₂ couple (−410 mV) and the NAD⁺/NADH couple (−320 mV) of the *C. necator* hydrogenase are closely aligned (Lauterbach et al. 2011a; 2011b). The directionality of H_2_ formation or dissociation can therefore be influenced through small perturbations in pH. Feeding sugars to *E. coli* results in the production of mixed acids that acidify the media (Sánchez-Clemente et al., 2018). Unlike sugars, oxidized substrates such as acetate do not produce organic acids and may instead alkalize the media by fixing protons during oxidative phosphorylation (Sánchez-Clemente et al., 2020). Increasing [H⁺] would favor H_2_ formation through substrate-product kinetics, which may explain why H_2_ consumption is difficult (this study) but H_2_ production is not (Lamont & Sargent, 2017; Ghosh et al., 2013; Schiffels et al., 2013).

Second, sugars such as glucose provide an abundant source of NAD(P)H through their metabolism via glycolysis and pentose phosphate pathways. Cellular NADH concentration is allosterically regulated through inhibition of TCA cycle enzymes and by transcriptional regulators (MacLean et al., 2023; Brown et al., 2022). Glucose cultivation would be expected to elevate NADH levels, which may decrease demand for exogenous reducing power provided by reversible hydrogenases.

In support of both hypotheses (they are not mutually exclusive), we reveal an early experiment demonstrating that H_2_ uptake by BW-HYD is low when cultivated on three fermentable substrates (glucose, xylose, and gluconate) and high when cultivated on four oxidized substrates (pyruvate, acetate, succinate, and malate; Figure S14 in *SI*).

### Economic Viability and Industrial Implementation

Using exogenous electron donors could improve the economics of biomanufacturing for commodity chemicals. While a thorough technoeconomic analysis (TEA) is needed, our data provide a basis for exploring its potential. We observed that up to 37.8% of the energy provided to *E. coli* through the oxidation of H_2_ can be used to enhance mevalonate formation (Table 1, Figure 4), thereby establishing an experimentally validated benchmark for efficiency. Glucose, the most widely used biomanufacturing feedstock, has a wholesale price of approximately $0.70 per kg in the US (March 2024: IMARC Group). To compensate for the oxidation of $1.00 worth of glucose would require the expenditure of 283 moles of H_2_. Excluding other technoeconomic considerations, if the price of hydrogen gas is $1.77 per kg or lower, the energy yield comparisons described in this study would suggest that a net reduction in operating expenses becomes feasible. If, through future metabolic engineering efforts, the percentage of oxidized H_2_ used to enhance bioproduction approaches 100% (similar to the 99.0 ± 2.8% efficiency of formate; Table 1, Figure 4), then only 107 moles of H_2_ would be required, raising H_2_ price parity to $4.67 per kg.

It is noteworthy that the price of non-renewable H_2_ is around $1.00 per kg (April, 2024: Business AnalytIQ), suggesting that price parity has already been achieved. Although non-renewable H_2_ will not provide systemic CO_2_ abatement throughout the production chain, it would allow industrialists to immediately develop H_2_-supported biomanufacturing until such time (or if) the cost of renewable H_2_ achieves price parity. The cost of electrolytic H_2_ is primarily determined by electricity prices (De Luna et al., 2019; Janssen et al., 2022), and as global energy capacity expands, the levelized cost of electricity from alternative sources is projected to decline by 50% by mid-century (Jayadev et al., 2020; Sens et al., 2022).

We demonstrated using formate-fed cells that chemical electron donors can be highly effective at conserving substrate (98.4 ± 3.6% for acetate; Table 1, Figure 3) and boosting product yield (99.0 ± 2.8% for mevalonate; Table 1, Figure 4), setting an optimistic benchmark for future biomanufacturing research. The immediate metabolic response seen in formate-fed cultures (Figure 3C) highlights a practical advantage of formate over hydrogen. While hydrogen uptake required a maturation period for enzyme activation, formate was rapidly absorbed, immediately replacing acetate oxidation and decreasing CO_2_ evolution. Although wholesale formic acid is similarly priced to H_2_, its low reductive capacity per unit mass (∼21.7 NADH/kg formate vs. ∼429 NADH/kg H_2_) makes cost parity with conventional feedstocks unlikely in the near term. Nonetheless, if cost parity can be achieved, formate holds great promise as an industrially appealing electron donor: It is soluble, easily crosses membranes, and directly supports NADH regeneration without enzymatic activation delays.

Economic trends suggest that H_2_ could serve as a viable alternative energy source, reducing reliance on feedstock-derived energy while enhancing bioprocess efficiency. From a process design perspective, the amount of money spent on feedstock just to produce the requisite biomass enabling commodity chemical production can be considerable, especially if batch schemes are used (Keasling et al. 2021). Moreover, whereas biomanufacturing ideally occurs within a steady state condition wherein carbon substrates are fed to microbial hosts and are completely converted into value-added chemicals, at least some portion of this feedstock is used for additional biomass accumulation. Our results show that both electron donors can support cellular energy needs (Figures 2 and 3) and enhance mevalonate biosynthesis (Figure 4) without spurring surplus cell growth. Conceptually, external reducing power could replace the reducing power ordinarily provided through the oxidation of feedstock, enabling greater bioreactor productivity per unit cost of raw materials. While we demonstrated this principle using mevalonate biosynthesis, the approach could be theoretically applicable to most bioproducts, as exemplified by our preliminary work on the NADH-independent flaviolin pathway (Figure S11 in *SI*).

### Broader Implications and Future Directions

The development of new technologies in transportation and manufacturing is crucial for American energy dominance and economic growth. Enhancing the efficiency of biomanufacturing through cost-effective chemical electron donors could play a key role in expanding global biomanufacturing capacity. Such a strategy could complement existing approaches, including the use of lignocellulosic biomass, gaseous waste streams, and cell-free biomanufacturing. The engineered strains described herein not only enable feedstock-efficient bioproduction but also contribute to a broader framework for the maturing biomanufacturing industry.

Future work should focus on optimizing hydrogenase expression stoichiometry, developing minimal maturase cassettes, and exploring the potential for creating NADH-intensive biosynthetic pathways that could improve H_2_ uptake on sugar substrates. Additionally, ^13^C-metabolic flux analysis would provide definitive validation of the glyoxylate shunt hypothesis and elucidate the precise metabolic rewiring that enables carbon conservation. These investigations will be essential for translating this proof-of-concept work into industrially viable bioprocesses that can compete economically with traditional petrochemical production. This technology should be translatable for most systems that require additional NADH, and may be exceptionally well-suited for cell-free biomanufacturing. We believe that this approach will be generally applicable to other industrially relevant organisms, and a similar approach has already been shown in *Pseudomonas putida* (Lonsdale et al., 2015). While the technology is promising, there are some practical limitations to deploying our technologies. Possible steps to derisking this technology include designing bioreactors that increase gas mass transfer rates while maintaining safe operation, formate-feeding schemes that minimize formate toxicity, and further organism engineering to maintain pathway redox balance without perturbing cellular fitness.

## CREDIT AUTHORSHIP CONTRIBUTION STATEMENT

**Robert L. Bertrand**: Conceptualization, Data curation, Formal analysis, Funding acquisition, Investigation, Methodology, Visualization, Writing – original draft. **Justin Panich**: Conceptualization, Data curation, Formal analysis, Funding acquisition, Investigation, Methodology, Visualization, Writing – original draft. **Aidan E. Cowan**: Methodology, Resources, Writing – Review and editing. **Jacob B. Roberts**: Methodology, Formal analysis, Writing – Review and editing. **Lesley J. Rodriguez:** Investigation, Writing – review and editing. **Juliana Artier**: Conceptualization, Writing – review and editing. **Emili Toppari**: Investigation, Writing – review and editing. **Edward E.K. Baidoo**: Investigation, Data curation, Formal analysis, Resources, Writing – review and editing. **Yan Chen**: Investigation, Data curation, Formal analysis, Resources, Writing – review and editing. **Christopher J. Petzold**: Investigation, Data curation, Formal analysis, Resources, Writing – review and editing. **Graham A. Hudson**: Investigation; Writing – review and editing. **Patrick M. Shih**: Conceptualization, Funding acquisition, Methodology, Project administration, Resources, Supervision, Writing – review and editing. **Steven W. Singer**: Conceptualization, Funding acquisition, Methodology, Project administration, Resources, Supervision, Writing – review and editing. **Jay D. Keasling**: Conceptualization, Funding acquisition, Methodology, Project administration, Resources, Supervision; Writing – review and editing.

## ACKNOWLEDGEMENTS

This work was funded by the Advanced Research Projects Agency-Energy (ARPA-E: *A Microbial Consortium Enabling Complete Feedstock Conversion*) and by the US Department of Energy (DOE) Joint BioEnergy Institute (https://www.jbei.org), supported by DOE, Office of Science, Biological and Environmental Research Program, under contract DEAC02-05CH11231 between DOE and Lawrence Berkeley National Laboratory. RLB acknowledges funding from the Natural Sciences and Engineering Research Council of Canada (NSERC: 532495-2019). JBR acknowledges support from a fellowship award under contract FA9550-21-F-0003 through the National Defense Science and Engineering Graduate (NDSEG) Fellowship Program, sponsored by the Air Force Research Laboratory (AFRL), the Office of Naval Research (ONR), and the Army Research Office (ARO). We thank Frank Sergeant (Newcastle University, United Kingdom) for genetic materials and Patrick Hallenbeck^†^ (Université de Montréal, Canada) for advice.

## DISCLAIMER

The United States Government retains, and the publisher, by accepting the article for publication, acknowledges that the United States Government retains a nonexclusive, paid-up, irrevocable, worldwide license to publish or reproduce the published form of this manuscript, or allow others to do so, for United States Government purposes. Any subjective views or opinions that might be expressed in the paper do not necessarily represent the views of the U.S. Department of Energy or the United States Government.

## AVAILABILITY OF SUPPORTING INFORMATION

Primers, plasmids, strains, supporting figures, and supporting tables are available in the accompanying *Supporting Information* file [LINK].

## AVAILABILITY OF GENETIC MATERIALS

Primers, plasmids, and strains are listed in Table S1-S3 of *SI* and are freely available for academic use via our institution’s Inventory of Composable Elements (ICE) (Ham et al., 2012). Please refer to the requisition numbers provided in the *SI* when submitting requests. Please inquire with our institution’s Intellectual Property Office regarding commercial use.

