## Supplemental Information for "Feedstock-Efficient Conversion through Hydrogen and Formate-Driven Metabolism in *Escherichia coli*"

### Table of Contents

*Supplemental Materials and Methods*:

Intracellular metabolite quantification 2
Protein identification and quantification 2
Mevalonate analysis by HPLC-MS 3
Computational flux balance analysis 4
Deductive metabolic estimates 4
Figure S1: Scheme – Construction of 13-gene hydrogenase plasmid 7
Figure S2: Scheme – End-point titration of NADH donors vs. $CO_2$ / acetate 8
Figure S3: Scheme – Time-course experiment titrating $H_2$, $CO_2$, and feedstock 8
Figure S4: Scheme – End-point assay of NADH donors vs. mevalonate yield 9
Table S1: Plasmids used in this study 9
Table S2: Primers used in this study 11
Table S3: Bacterial strains used in this study 12
Table S4: Statistical variation in gas pipetting technique 13
Table S5: Statistical variation in statistical replicate cultures 14
Table S6: Statistical variation in gas analysis 15
Table S7: Statistical variation in HPLC analysis 16

*Supplemental Results and Discussion:*

Figure S5: Enlarged plasmid map of HYD (Kan | P15A) 17
Figure S6: Effects of $H_2$ on intracellular TCA metabolite concentration 18
Figure S7: $H_2$ uptake measurements pertinent to Figure 2 19
Figure S8: Linear correlations between substrates, CO2 evolution, feedstock 20
Figure S9: Oxygen gas consumption metrics pertinent to Figure 2C and 3C 21
Figure S10: Formate uptake and total $CO_2$ measurements pertinent to Figure 3 21
Figure S11: Enhanced flaviolin production by $H_2$ 22
Figure S12: Mevalonate titer achieved when using acetate-based media 23
Figure S13: Enhancement of $H_2$ uptake by pre-incubation method 24
Table S8: Molar quantities of mixed acid products from Figure 4 25
Figure S14: $H_2$ uptake by *E. coli* various sugars and oxidized substrates 26

*Supplemental References*

References 27

**Intracellular metabolite quantification:** The intracellular metabolite profile was evaluated using LC-MS. Samples were prepared by cooling 1.5 ml broth aliquots in an ice slurry, followed by centrifugation to remove the supernatant. To extract metabolites and quench reactions, the pellet was resuspended in 250 µL of cold methanol, followed by 250 µL of cold water. To remove enzymes, the resuspension was centrifuged, and the resultant supernatant was filtered through an Amicon 3 kDa MWCO spin column (*Millipore*). The filtrate was diluted to 1 ml with water, snap-frozen with liquid nitrogen, and lyophilized at -105°C for 36h (*Labconco*). Dry metabolites were resuspended in 100 µL of 50% methanol in water and stored at -80°C until needed. Organic acids were measured via reversed-phase chromatography and high-resolution mass spectrometry. Liquid chromatography – mass chromatography (LC-MS) was performed as previously described (Wang et al., 2025), as follows: LC was conducted on an Ascentis Express RP-Amide column (150-mm length, 4.6-mm internal diameter, and 2.7-µm particle size) (*Sigma-Aldrich*), equipped with the appropriate guard column, using an Agilent Technologies 1260 Series high-performance liquid chromatography (HPLC) system (*Agilent*). A sample injection volume of 2 µL was used throughout. The sample tray and column compartment were set to 5 and 50°C, respectively. The LC-MS-grade mobile phase solvents were purchased from *Honeywell*, and all other reagents were purchased from *Sigma-Aldrich*. The mobile phases were composed of 0.1% formic acid/84.9% water/15% methanol (v/v/v) (A), 0.04% formic acid and 5 mM ammonium acetate in methanol (B). Metabolites were separated via gradient elution under the following conditions: Linearly increased from 0% B to 30% B in 5.5 min, increased to 100% B in 0.2 min, held at 100% B for 2 min, decreased from 100% B to 0% B in 0.2 min, and held at 0% B for 2.5 min. The flow rate was held at 0.4 ml/min for 7.7 min, linearly increased from 0.4 ml/min to 1 ml/min in 0.2 min, and held at 1 ml/min for 2.5 min. The total LC run time was 10.4 min. The HPLC system was coupled to an Agilent Technologies 6545 series quadrupole time-of-flight mass spectrometer (QTOF-MS) and conducted as follows: Drying and nebulizing gasses were set to 10 L/min and 25 psi, respectively, and drying gas temperature of 300°C was used throughout. Sheath gas temperature and flow rate were 330°C and 12 L/min, respectively. Electrospray ionization via the Agilent Technologies Jet Stream Source was conducted in the negative ion mode using a capillary voltage of 3,500 V. The fragmentor, skimmer, and OCT 1 RF Vpp voltages were set to 100, 50, and 300 V, respectively. The acquisition range was from 70-1,100 m/z, and the acquisition rate was 1 spectra/sec. Prior to data acquisition, the QTOF-MS system was tuned with the Agilent ESI-L Low concentration tuning mix (diluted 10-fold in a solvent mixture of 80% acetonitrile and 20% water) in the range of 50-1700 m/z. Reference lock masses from data acquisition and processing were conducted via the Agilent MassHunter software package. Reference mass correction was performed with 5 µM trifluoroacetic acid ammonium salt (part number I8720243) and 5 µM HP-0921 (part number 18720241) at a flow rate of 5 µL/min via a second ESI sprayer. Data processing and analysis were conducted via Agilent MassHunter Qualitative Analysis, Agilent Profinder, and/or Agilent MassHunter Quantitative Analysis. All other metabolites were analyzed according to the reported methods (Amer et al., 2022) and quantified via seven-point calibration curves ranging from 0.39 to 25 µM.

**Protein identification and quantification:** Analysis and quantification of protein content were performed by mass spectrometry proteomics. Five milliliters of bacterial cultures were centrifuged, and the supernatant was discarded. The pellet was twice washed with PBS and stored at -80 °C until needed. Protein was extracted using an established proteomic sample preparation protocol (Chen et al., 2023), as follows: Cell pellets were resuspended in P2 Lysis Buffer (*Qiagen*). Proteins were precipitated with the addition of 1 mM NaCl and four volumes of acetone, followed by two additional washes with 80% acetone in water. The recovered protein pellet was homogenized by pipetting and mixed with 100 mM ammonium bicarbonate in 20% methanol. Protein concentration was determined by the DC protein assay (*BioRad*). Protein reduction was accomplished using 5 mM tris 2-(carboxyethyl)phosphine (TCEP) for 30 min at room temperature, and alkylation was performed with 10 mM iodoacetamide (IAM; final concentration) for 30 min at RT in darkness. Overnight digestion with trypsin was accomplished with a 1:50 trypsin:total protein ratio. The resulting peptide samples were analyzed on an Agilent 1290 UHPLC system coupled to a ThermoScientific Orbitrap Exploris 480 mass spectrometer. Peptide samples were loaded onto an Ascentis ES-C18 Column (*Sigma–Aldrich*) and were eluted from the column by using a 10-minute gradient from 98% solvent A (0.1 % formic acid in water) and 2% solvent B (0.1% formic acid in acetonitrile) to 65% solvent A and 35% solvent B. Eluting peptides were introduced to the mass spectrometer operating in positive-ion mode and were measured in data-independent acquisition (DIA) mode with a duty cycle of 3 survey scans from m/z 380 to m/z 985 and 45 MS2 scans with precursor isolation width of 13.5 m/z to cover the mass range. DIA raw data files were analyzed by an integrated software suite DIA-NN. The database used in the DIA-NN search (library-free mode) is the *E. coli* latest Uniprot proteome FASTA sequences plus the protein sequences of the heterologous proteins and common proteomic contaminants. DIA-NN determines mass tolerances automatically based on first-pass analysis of the samples with automated determination of optimal mass accuracies. The retention time extraction window was determined individually for all MS runs analyzed via the automated optimization procedure implemented in DIA-NN. Protein inference was enabled, and the quantification strategy was set to Robust LC = High Accuracy. Output main DIA-NN reports were filtered with a global FDR = 0.01 on both the precursor level and protein group level. The Top3 method, which is the average MS signal response of the three most intense tryptic peptides of each identified protein, was used to plot the quantity of the targeted proteins in the samples (Ahrné et al., 2013; Silva et al., 2006). The generated mass spectrometry proteomics data was deposited to the ProteomeXchange Consortium via the PRIDE partner repository, with dataset identifier PXD061595 (Perez-Riverol et al., 2022).

**Mevalonate analysis by LC-MS:** Mevalonate produced by BW-HYD-MEV and CM-HYD-MEV when cultivated in EZ-Acetate was quantified by reverse phase, high-performance liquid chromatography and mass spectrometry (RP-HPLC-MS). Samples were prepared by isolating broth supernatant followed by the addition of 10% methanol to create an internal loading standard. Samples were filtered using 0.45 μm modified nylon centrifugal filters (*VWR*), and filtrates were stored at -80°C until needed. Analysis was performed using an Agilent 1260 Infinity II HPLC with an MSD/iQ mass spectrometry detector equipped with an analytical EC UHPLC Nucleodur C18 Htec 1.8 μm 100×2 mm column (Macherey Nagel, 760306.20). Analysis occurred at room temperature, with 10 μl injection volume, using LC-MS grade water containing 0.1% formic acid (buffer A) and LC-MS grade acetonitrile containing 0.1% formic acid (buffer B), via the following gradient: 0 min 2% B, 4 min 7.5% B, 5 min 95% B, and 6min 2% B, with a 14-minute hold for equilibration between injections. Mevalonate was monitored by a negative mode single-ion monitoring (SIM) at m/z = -147.1 and exhibited a retention time of 2.548 minutes. Mevalonate was interpolated using a standard plot (1 to 100 mg/L) of an authenticated standard. Mevalonate produced by these strains when cultivated in M9-Glucose was quantified by HPLC, as described in the *Main Text*.

**Computational Flux Balance Analysis:** The marginal yield of various products on hydrogen gas and formate was calculated in Python using the COBRApy flux balance analysis software (Ebrahim et al., 2013). The curated metabolic model for *E. coli*'s core carbon metabolism was downloaded from BIGG (King et al., 2016; Orth et al., 2010 (Wenk et al., 2020). Yield envelopes are typically reported as the entire set of possible production yields of multiple outputs, commonly used to model the possible tradeoffs between biomass production and the production of some product (Klamt et al., 2018). We introduce the term "marginal yield envelope" to mean the yield envelope of a system with a small perturbation in its inputs, which had been growing at its maximal rate. The marginal yield envelope was determined by calculating the maximal change in each product's titer via incremental changes in the feedstock. We assume that the system's response to incremental perturbations is linear, and that sufficiently small additions of hydrogen gas or formate to a system that is growing at a maximal growth rate will result in small and proportional changes in the system's product formation rate. This linear response assumption is justified by the continuity and differentiability of the underlying biological functions modeled in our system. For example, to calculate the marginal yield of acetate on hydrogen gas: (1) Load the curated *E. coli* core model (Wenk et al., 2020), add the hydrogenase net reaction, and include synthesis pathways; then (2) Calculate the maximal growth rate, which is used as a lower bound for the following two calculations; (3a) Calculate the maximum acetate production flux of the system without hydrogen gas influx, defined as $F_{ACETATE\ -\ NO\ H2}$ in mmol/gCDW/hr; (3b) Calculate the maximum acetate production flux of the system with a hydrogen gas influx upper bound of 1.0 mmol/gCDW/hr, defined as $F_{ACETATE\ YES\ H2}$ in mmol/gCDW/hr; (4) Calculate marginal yield (mol acetate / mol hydrogen gas):

$$Marginal\ Yield_{ACETATE\ H2} = \frac{\Delta F_{ACETATE}}{\Delta F_{H2}} = \frac{F_{ACETATE\ YES\ H2} - F_{ACETATE\ NO\ H2}}{1.0\ mmol\ H_2/gCDW/hr}$$

This process is repeated for all other products and electron donors.

**Manual Metabolic Estimates:** A manual estimate of the bioenergy requirements for each cellular event was generated to sanity check computational FBA. As shown below, this can be achieved once necessary approximations are permitted (Milo & Phillips, 2015).

*Using proton motive force equivalencies*: The key approximation enabling the manual estimation of bioenergy requirements is equating ATP, $FADH_2$, NADH, and NADPH to proton motive force (PMF). While the electrons donated by NADH to the human respiratory chain are well known to power the translocation of ten protons ($e^-/H^+ = 5$), this ratio in *E. coli varies* due to its facultative aerobe nature. Among respiring *E. coli*, the $e^-/H^+$ is 4 but can decrease during anaerobic cultivation (Sharma et al., 2012; Borisov et al., 2011). As our *E. coli* cultures in butyl-stoppered serum vials are initiated in air and observed to respire throughout an overnight cultivation (Figure

2C, *main text*), we estimated that the $e^-/H^+$ of NADH is 4. Consequently, we equated NADH in *E. coli* to 8 $H^+$ in PMF. To our knowledge, the $e^-/H^+$ of $FADH_2$ in *E. coli* is not well established. In the absence of information, we deferred to the human $e^-/H^+$ ratio of 3, equating $FADH_2$ to 6 $H^+$ in PMF. It is well established in *E. coli* and other bacteria that four $H^+$ in PMF are expended to phosphorylate ADP to ATP (Steigmiller et al., 2008; ). Thus, each ATP is equated to 4 $H^+$ in PMF. Finally, hydride exchange from NADH to $NADP^+$ is possible in *E. coli* via the enzyme PntAB but requires proton translocation (Sauer et al., 1999). Therefore, each NADPH is equated to 9 $H^+$ in PMF.

*Estimating the energy provided by hydrogen gas and formic acid*: All experiments were conducted in butyl-stoppered serum vials. The transfer efficiency of hydride equivalents from $H_2$ to $NAD^+$ by hydrogenase exceeds 99% in ambient air (Lauterbach & Lenz, 2013). Thus, the oxidation of each $H_2$ is equated to the donation of 8 $H^+$ in PMF. The passive diffusion of formic acid across the hydrophobic cell membrane is greatly favored over the charged formate species. Any protons that enter the cell also contribute to the electrochemical potential. When reducing power is supplied as formate (such as in this work), it must first recruit a proton from the broth before entering the cell. Therefore, the oxidation of each formic acid is equated to the donation of 9 $H^+$ in PMF.

*Estimating the energy required to prevent the complete oxidation of acetate*: To catabolize acetate, *E. coli* must first activate it to acetyl-CoA, at the expense of ATP. The irreversible acetyl-CoA synthetase pathway predominates when *E. coli* is cultivated in defined media with acetate as the primary carbon source and when exposed to declining oxygen partial pressure (Kumari et al., 2000). Therefore, acetate consumption requires an investment of two ATP. Subsequently, acetyl-CoA is oxidized to $CO_2$ by the Krebs cycle, yielding 3 NADH (3 x 8 $H^+$), 1 $FADH_2$ (1 x 6 $H^+$), and 1 ATP (1 x 4 $H^+$). Thus, acetate provides net cellular energy equivalent to 26 $H^+$ in PMF. To prevent the oxidation of one mole of acetate, we estimate that 3.25 (26/8) moles of $H_2$ or 2.89 (26/9) moles of formic acid are required.

*Estimating the energy required to form surplus mevalonate from glucose*: The decarboxylative glycolysis of glucose ends with acetyl-CoA. The formation of mevalonate requires three acetyl-CoA and two NADH. Therefore, the oxidation of acetyl-CoA must be suppressed to enable its conversion into surplus mevalonate. The digestion of acetyl-CoA produces three NADH (3 x 8 $H^+$), one $FADH_2$ (1 x 6 $H^+$), and 1 ATP (1 x 4 $H^+$), totaling 34 $H^+$ in PMF. Consequently, three acetyl-CoA molecules require 102 $H^+$ in PMF. Additionally, two NADH (2 x 8 $H^+$) are necessary, bringing the total to 118 H+. Thus, to facilitate the formation of one additional mole of mevalonate, we estimate that 14.75 (118/8) moles of $H_2$ or 13.11 (118/9) moles of formic acid are needed.

*Estimating the energy required to ferment surplus acetate from glucose*: By fermenting acetate from acetyl-CoA, *E. coli* forfeits the cellular energy it would have gained by oxidizing it, minus the one ATP it recuperated via the PTA-ACKA pathway. Therefore, the loss of 3 NADH (3 x 8 $H^+$), 1 $FADH_2$ (1 x 6 $H^+$), and 1 ATP (1 x 4 $H^+$), contrasted with the gain of 1 ATP (1 x 4 $H^+$), results in a net energy implication of 30 $H^+$ PMF. Thus, to increase acetate fermentation by one mole, we estimate that 3.75 (30/8) moles of $H_2$ or 3.33 (30/9) moles of formic acid are required.

*Estimating the energy required to ferment surplus ethanol from glucose*: By fermenting ethanol, *E. coli* forfeits the cellular energy it could have gained by oxidizing acetyl-CoA and discards two additional NADH to reduce acetyl-CoA to ethanol via the acetaldehyde intermediate. This equals 5 NADH (5 x 8 $H^+$), 1 $FADH_2$ (1 x 6 $H^+$), and 1 ATP (1 x 4 $H^+$), or 50 $H^+$ PMF. Therefore, to enhance ethanol fermentation by one mole, we estimate that 6.25 (50/8) moles of $H_2$ or 5.56 (50/9) moles of formic acid are needed.

*Estimating energy required to ferment surplus formate from glucose*: Formic acid is fermented through the non-oxidative decarboxylation of pyruvate into acetyl-CoA via pyruvate formate lyase, which results in the loss of one NADH (8 $H^+$ in PMF) that would have been produced if pyruvate were instead oxidatively decarboxylated into acetyl-CoA via the pyruvate dehydrogenase pathway. Subsequently, the release of formic acid into the extracellular space transports a proton from the intermembrane space, which decreases the electrochemical potential. Therefore, to increase formate fermentation by one mole, we estimate that 1.125 (9/8) moles of $H_2$ or 1.00 (9/9) moles of formic acid are needed.

*Estimating energy required to increase E. coli dry biomass*: By analogy, building a house incurs financial costs for both parts and labor. We must therefore estimate the amount of cellular energy needed to prevent the oxidation of glucose into $CO_2$ ("parts") as well as the energy required to turn glucose into biomass ("labor"). The complete oxidation of one mole of glucose (180.156g) yields 10 NADH (80 $H^+$), 2 $FADH_2$ (12 $H^+$), and 4 ATP (16 $H^+$), for a total of 108 moles $H^+$ in PMF. The "parts" required to build 180.156g of dry *E. coli* biomass is therefore estimated to be 108 moles $H^+$ in PMF, or 0.599 moles $H^+$ in PMF per gram of dry biomass. The ATP consumption rate of exponentially growing *E. coli* was measured at 6,400,000 ATP molecules (25,600,000 $H^+$ in PMF) per cell per second (Deng et al., 2021). The weight of an exponentially growing *E. coli* bacterium is approximately 200 femtograms (Loferer-Krößbacher et al., 1998; Heldal et al., 1985). Assuming it takes 30 minutes for one gram of dry biomass to double, the "labor" portion is estimated to equate to 0.383 moles $H^+$ in PMF per gram of dry biomass. The combination of "parts" and "labor" is 0.982 moles $H^+$ in PMF per gram of dry *E. coli* biomass. Therefore, to increase dry biomass by one gram, we estimate that 0.123 (0.982/8) moles of $H_2$ or 0.109 (0.982/9) moles of formic acid are required.

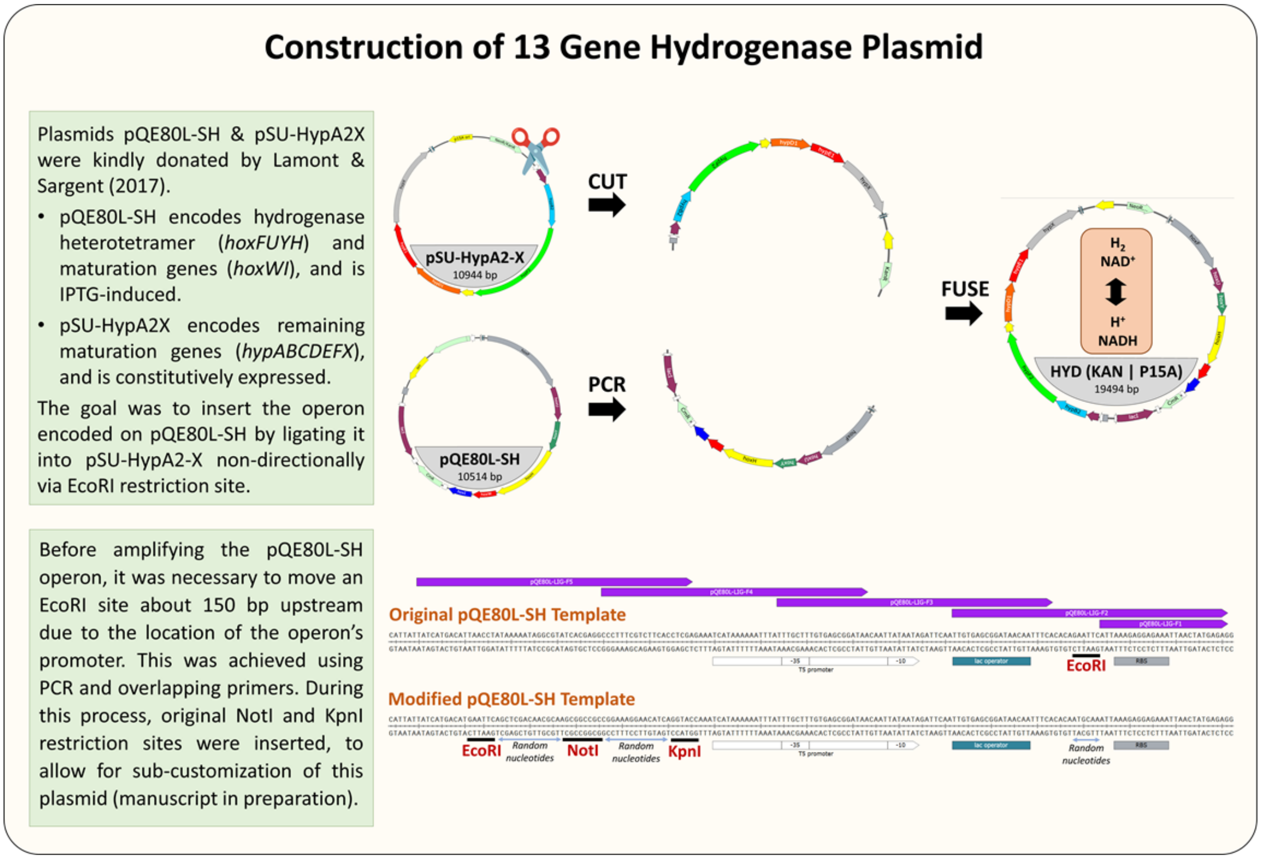


**Figure S1: Scheme for constructing a single plasmid that encodes the *C. necator* hydrogenase heterotetramer and nine maturation proteins for heterologous expression in *E. coli*.** Refer to *DNA manipulations* and *Prototyping the C. necator soluble hydrogenase (HYD) and Pseudomonas sp. 101 formate dehydrogenase (FDH)* in the *Main Text*.

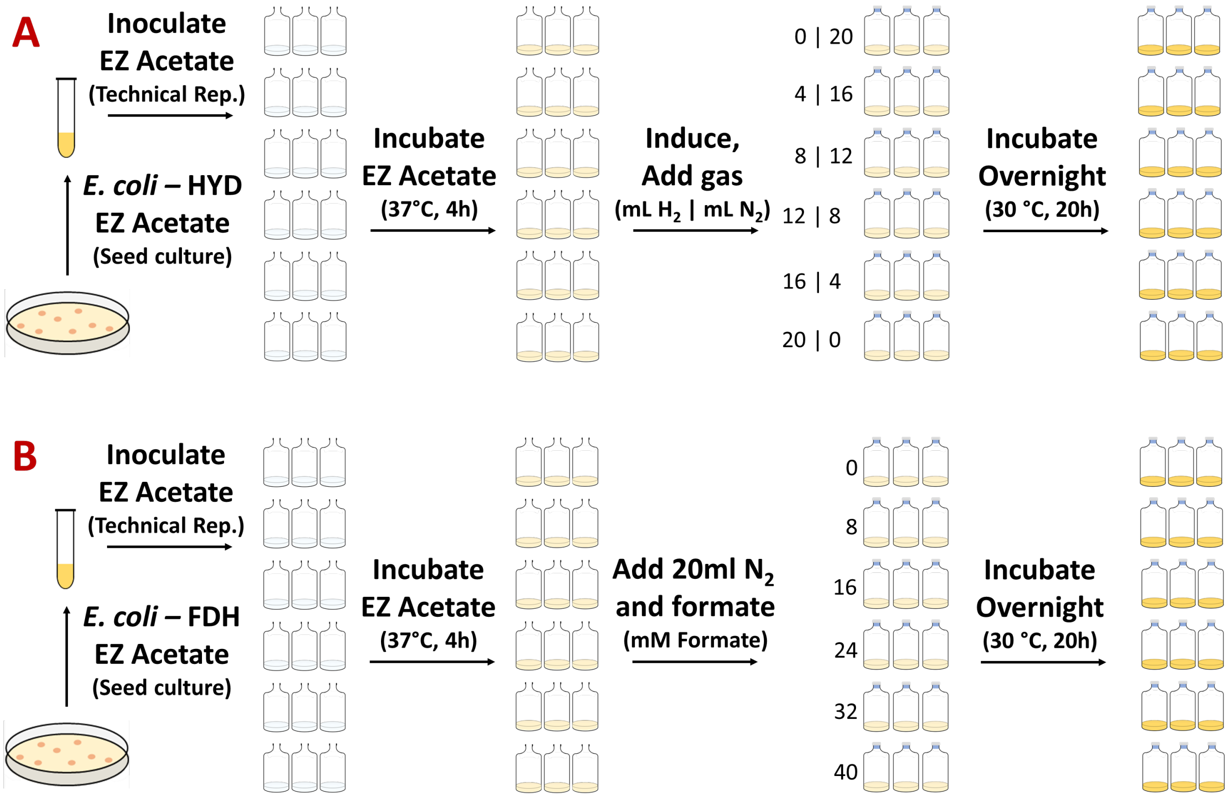


**Figure S2: Scheme of end-point titration to observe the effects of hydrogen gas (A) and formate (B) on acetate consumption and $CO_2$ production in BW-HYD and BW-FDH.** Refer to *Gas cultivations* and *Electron donors, which prevent the decarboxylation of an organic carbon feedstock* in the *Main Text*.

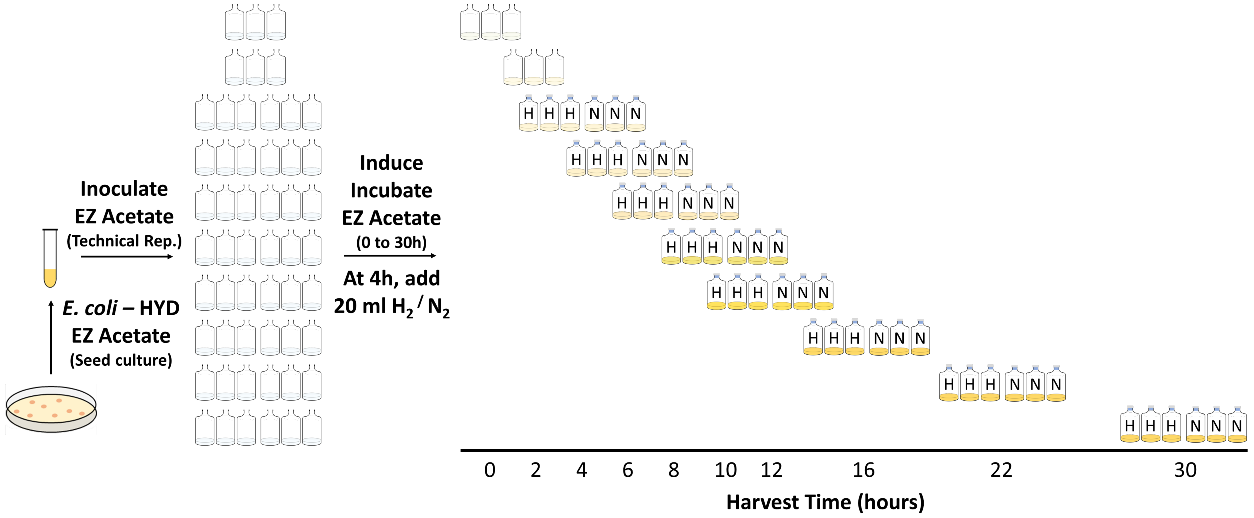


**Figure S3: Scheme of a time-course titration to observe the effects of hydrogen gas on acetate consumption and $CO_2$ production in BW-HYD.** Refer to *Gas cultivations* and *Electron donors, which prevent the decarboxylation of an organic carbon feedstock* in the *Main Text*.

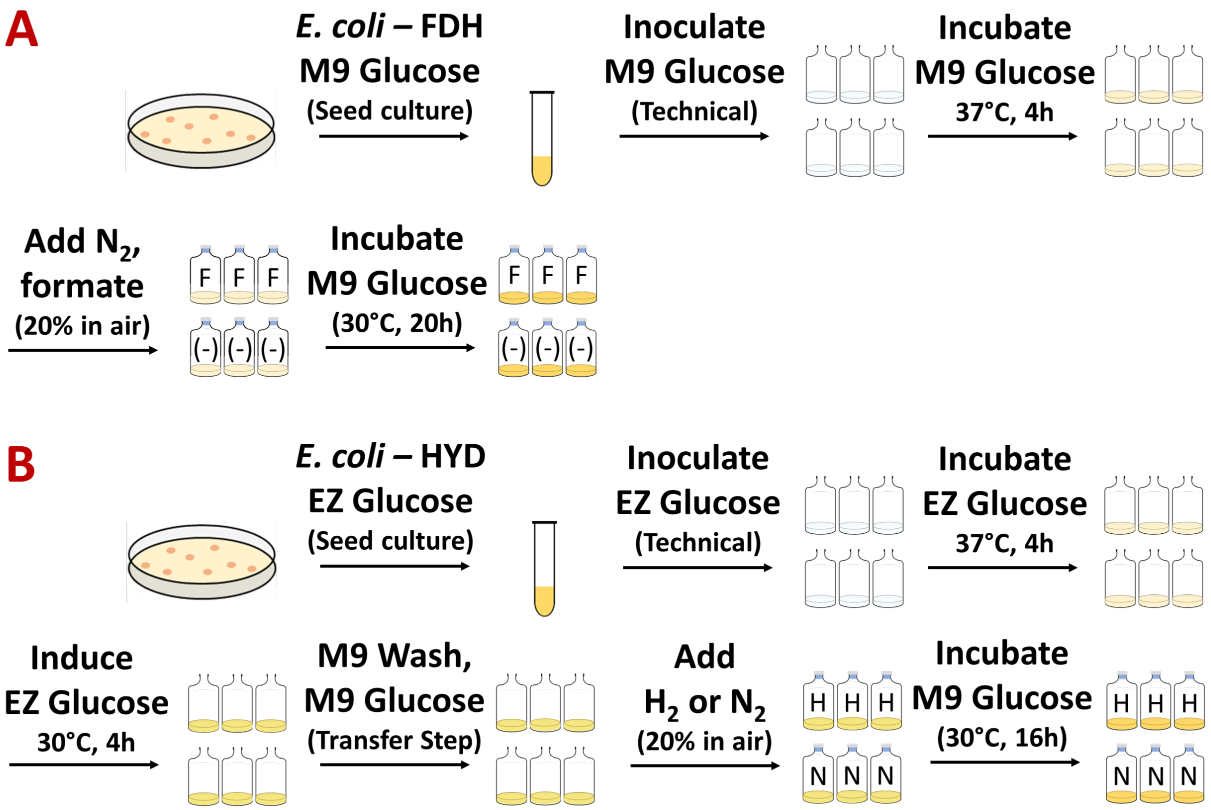


**Figure S4**: **Scheme for assessing the effects of formate on mevalonate production in (A) BW-FDH-MEV and CM-FDH-MEV, and (B) hydrogen gas on BW-HYD-MEV and CM-HYD-MEV. The u**ptake of $H_2$ gas is limited when *E. coli* is cultivated in minimal media. Greater $H_2$ uptake was achieved by pre-cultivating *E. coli* in rich media and then transferring it to minimal media. Refer to '*Gas cultivations* and *Electron donors increase mevalonate yield with optimizable efficiency*' in the *Main Text*.

**Table S1: Plasmids constructed or used in this study.**

| **Name** | **Description** | **ICE Part No.** | **Reference** |
|---|---|---|---|
| pQE80L-SH | Encodes hydrogenase heterotetramer (*hoxFUYH*) and maturation genes (*hoxWI*). Used to build HYD (Kan:P15A) (See Figure S1). Uses IPTG induction, carbenicillin resistance, and ColE1 origin. 10.5 KB. | Please request from original authors | Lamont & Sargent, 2017 |
| pSU-HypA2X | Encodes hydrogenase maturation genes (*hypABCDEFX*). Used to build HYD (Kan:P15A) (See Figure S1). Uses constitutive expression, kanamycin resistance, and P15A origin. 10.9 KB. | Please request from original authors | Lamont & Sargent, 2017 |
| HYD (Kan:P15A) | Encodes hydrogenase heterotetramer and nine material genes. The *hoxFUYHWI* operon is IPTG | Jbx_258372 | This study |

| | | | |
|---|---|---|---|
| | inducible. The *hypABCDEFX* operon is constitutively expressed. Contains a non-functional chloramphenicol marker as an artifact. Uses kanamycin resistance and P15A origin. 19.4 KB. | | |
| HYD (Spec: P15A) | A derivative hydrogenase plasmid that is otherwise identical to HYD (Kan:P15A) but instead uses a spectinomycin selection marker, constructed using a derivatization assembly protocol (manuscript in preparation). 19.0 KB. | Jbx_258374 | Manuscript forthcoming |
| FDH (Spec: ColE1) | Encodes NADH-specific formate dehydrogenase. Uses constitutive expression, spectinomycin resistance, and ColE1 origin. 3.3 KB. | Please request from original authors | Gleizer et al., 2019 |
| MEV (Chlor: SC101) | Low copy variant to promote cell growth used for hydrogen gas assays. Encodes acetoacetyl-CoA thiolase, HMG-CoA synthase, HMG-CoA reductase. Uses IPTG induction, chloramphenicol resistance, and SC101 origin. 9.0 KB. | JBx_258644 | This study |
| MEV (Chlor: P15A) | Medium copy producer of mevalonate used for formate assays. Encodes acetoacetyl-CoA thiolase, HMG-CoA synthase, HMG-CoA reductase. Uses IPTG induction, chloramphenicol resistance, and P15A origin. 7.5 KB. | JBx_258643 | Kang et al., 2019 |
| MEV (Chlor: ColE1) | High copy producer of mevalonate that was abandoned during this study due to deleterious effects on *cell* growth (data not shown). Uses IPTG induction, chloramphenicol resistance, and ColE1 origin. 7.9 KB. | JBx_258642 | This study |
| RppA | Encodes flaviolin type III polyketide synthase. Uses arabinose induction, chloramphenicol resistance, and BBR1 origin. 7.2 KB. | JBx_258390 | Incha et al. 2020 |
| HYD[A] (Hall) | An alternative hydrogenase construct not used in this study; Encodes *C. necator* HoxFUYHWI and HypA2B2R2. Codon and RBS composition are native to *C. necator*; BBR1 origin and arabinose induction. 13.3 KB. | JBx_258645 | Ghosh et al., 2013 |

[A]During this work, we re-created the hydrogenase plasmid originally reported by Ghosh et al., 2013, with advice provided by Patrick Hallenbeck. Preliminary evidence suggests that this re-creation is functional (data not shown). To our knowledge, this plasmid is the only surviving copy from the laboratory of this late professor and hydrogenase pioneer. This plasmid is available via our institution's ICE registry.

**Table S2: Primers used in this study**

| Name | Purpose | Sequence (5' – 3') |
|---|---|---|
| QE80-Lig-F1 | Assemble HYD (see Fig. S1) | ATTAAAGAGGAGAAATTAACTATGAGAG |
| QE80-Lig-F2 | Assemble HYD (see Fig. S1) | TTGTGAGCGGATAACAATTTCACACAATGCAAATTAAAGAGGAGAAATTAACTATGAGAG |
| QE80-Lig-F3 | Assemble HYD (see Fig. S1) | TTTGCTTTGTGAGCGGATAACAATTATAATAGATTCAATTGTGAGCGGATAACAATTTCA |
| QE80-Lig-F4 | Assemble HYD (see Fig. S1) | CGGAAAGGAACATCAGGTACCAAATCATAAAAAATTTATTTGCTTTGTGAGCGGATAA |
| QE80-Lig-F5 | Assemble HYD (see Fig. S1) | CGACGGCCAGTGAATTCAGCTCGACAACGCAAGCGGCCGCCGGAAAGGAACATCAGGTAC |
| QE80-Lig-R | Assemble HYD (see Fig. S1) | TTGCGCCAACCGACAGAATTCCCTGATGCGGTATTTTCTCC |
| Mev-ColE1F | Convert MEV-P15A into MEV-ColE1 | GCCCGCGCCTAATGAGGATCCAAACTCGAGCACATTTCCCCGAAAAGTGCC |
| Mev-ColE1R | Convert MEV-P15A into MEV-ColE1 | TTGAAGAGATAAATTGCACTGAAATCTAGAATTTCGATAAGCCAGCTGGGC |
| Mev-SC101F | Convert MEV-P15A into MEV-SC101 | GCCCGCGCCTAATGAGGATCCAAACTCGAGCCAGGCATCAAATAAAACGAAAGGC |
| Mev-SC101R | Convert MEV-P15A into MEV-SC101 | TTGAAGAGATAAATTGCACTGAAATCTAGAACTAGGGACAGTAAGACGGGTAAG |

**Table S3: Strains used in this study**

| Name | Description | ICE Strain No. | Reference |
|---|---|---|---|
| BW | Untransformed BW25113 *E. coli* (with DE3 complement enabling use of T7 Promoters). | JBx_258683 | This study |
| CM | Untransformed CM15 *E. coli* containing mutations knocking out major mixed acid production pathways (ΔaraBAD ΔFadRE ΔldhA ΔackApta ΔadhE ΔpoxB ΔfrdABCD) | Please request from original inventors | Mehrer et al., 2018 |
| BW-HYD | BW25113 *E. coli* containing IPTG-inducible HYD (Kan:P15A) plasmid (Jbx_258372) | Jbx_258690 | This study |
| BW-FDH | BW25113 *E. coli* containing constitutively expressed FDH (Spec:ColE1) plasmid (N/A) | Please request "BW" from us and plasmid from original inventors | Gleizer et al., 2019; This study |
| BW-HYD-MEV | BW25113 *E. coli* containing IPTG-HYD (Spec:P15A) plasmid (Jbx_258374) and IPTG-MEV (Chlor:SC101) plasmid (Jbx_258394) | Jbx_258701 | Kang et al., 2019; This study |
| BW-FDH-MEV | BW25113 *E. coli* containing constitutive FDH (Spec:ColE1) plasmid (N/A) and IPTG-inducible MEV (Chlor:P15A) plasmid (JBx_258644) | Please request "BW" and "MEV" from us, and "FDH" from original inventors | Gleizer et al., 2019; This study |
| BW-HYD-RppA | BW25113 *E. coli* containing IPTG HYD (Kan:P15A) plasmid (Jbx_258372) and arabinose-inducible RppA (Chlor:BBR1) plasmid (JBx_258390) | Jbx_258702 | Incha et al., 2020; this study |

| | | | |
|---|---|---|---|
| CM-HYD-MEV | CM15 *E. coli* containing IPTG HYD (Spec:P15A) plasmid (Jbx_258374) and IPTG MEV (Chlor:SC101) plasmid (JBx_258644) | Please request plasmids from us, and "CM" from original inventors | Mehrer et al., 2018; Kang et al., 2019; This study |
| CM-FDH-MEV | CM15 *E. coli* containing constitutive FDH (Spec:ColE1) plasmid (N/A) and IPTG-inducible MEV (Chlor:P15A) plasmid (JBx_258643) | Please request "MEV" from us, and "FDH" and "CM" from original inventors | Mehrer et al., 2018; Gleizer et al., 2019; This study |

**Table S4: Statistical variation in gas pipetting technique.** To evaluate the reproducibility of hydrogen gas injections, mock cultures were created using the same procedure typically used for preparing live cultures (see Materials and Methods in *Main Text*) as follows: Six serum bottles containing 10 ml of water were incubated with loose-fitting aluminum foil covers for 4h at 37°C and 150 RPM. The vials were then moved to a 30°C incubator and sealed with butyl stoppers. Using a syringe with a stopcock, 20 ml of air was first withdrawn from each serum vial, followed by the injection of 20 ml of $H_2$ into each vial. The vials each contained 10 ml of water and approximately 112 ml of total headspace. The six vials filled with water and gasses were incubated overnight at 30°C and 150 RPM. The following day, the gas composition of each serum vial was assessed using gas chromatography. The molar density of $H_2$ was defined as 39.95 µmol/ml at standard pressure and 30°C, and the consensus signal intensity is defined as exactly 20 ml of $H_2$ (799 µmol). The outcome of this control experiment indicated that 799 µmol of $H_2$ (20 ml) is injectable into serum vials with a margin of error of ± 5 µmol (0.14 ml).

| Replicate Number | Peak Area (unitless) | H2 (ml) | H2 (µmol) |
|---|---|---|---|
| 1 | 3851296 | 20.11 | 808 |
| 2 | 3795698 | 19.92 | 796 |
| 3 | 3826904 | 20.09 | 802 |
| 4 | 3814848 | 20.02 | 799 |
| 5 | 3780454 | 19.84 | 792 |
| 6 | 3793533 | 19.91 | 795 |
| **Average** | 3810456 | 20.00 | 799[a] |
| **Std. Dev.** | 25931 | 0.14 | 5 |
| **Error (%)** | 0.68 | 0.68 | 0.68 |

[a]In *Main Text* and *SI*, 799 µmol $H_2$ is rounded to 800 µmol for simplicity.

**Table S5: Statistical variation in replicate cultures**. To evaluate the reproducibility of live cultures, control cultures were prepared using the same procedure (see Materials and Methods in *Main Text*), as follows: A glycerol stock of BW25113-DE3 transformed with HYD ("BW-HYD") was plated on kanamycin agar and incubated overnight at 37°C. A single colony was then inoculated into 10 ml of EZ Rich-Acetate (55.5mM) and incubated aerobically overnight at 37°C and 200 RPM. A single seed culture was then used to inoculate six serum vials, each containing 10 ml of EZ Rich-Acetate (55.5 mM). Cultures were incubated for 4h at 37°C and 150 RPM. These cultures were subsequently transferred to a 30°C incubator and butyl-stoppered. Using a syringe with a stopcock, 20 ml of air was first withdrawn from each serum vial, followed by an injection of 20 ml of inert $N_2$. The cultures were then incubated overnight at 30°C and 150 RPM. On the following day, optical density (OD600), dry cell weight (g/L), $CO_2$ production (μmol), and acetate remaining in broth (μmol) were evaluated. Results reveal that this cultivation protocol introduces an error of 1-4% among statistical replicate cultures concerning these biochemical metrics.

| **Replicate Number** | **Density (OD600)** | **DCW (g/L)** | **$CO_2$ (μmol)** | **Acetate (μmol)** |
|---|---|---|---|---|
| 1 | 3.16 | 0.82 | 845 | 143 |
| 2 | 3.09 | 0.91 | 868 | 150 |
| 3 | 3.18 | 0.87 | 850 | 136 |
| 4 | 3.18 | 0.90 | 842 | 139 |
| 5 | 3.12 | 0.87 | 868 | 145 |
| 6 | 3.20 | 0.88 | 848 | 141 |
| **Average** | 3.16 | 0.87 | 854 | 142 |
| **Std. Dev.** | 0.04 | 0.03 | 11.7 | 4.8 |
| **Error (%)** | 1.33 | 3.56 | 1.37 | 3.37 |

**Table S6: Statistical variation in gas analysis.** To evaluate the intrinsic error of the gas chromatography unit (GC2014, *Shimadzu*), six syringes with stopcocks were assembled. To prevent contamination of gas samples by human breath, six gas aliquots were collected from an offline anaerobic growth chamber and then injected into the GC. The statistical deviation of the $N_2$ signal intensity was defined as the intrinsic error of the analytical unit.

| Replicate Number | Peak Area (unitless) |
|---|---|
| 1 | 1624904 |
| 2 | 1621471 |
| 3 | 1618126 |
| 4 | 1615792 |
| 5 | 1617353 |
| 6 | 1617199 |
| **Average** | 1619141 |
| **Std. Dev.** | 3402 |
| **Error (%)** | 0.21 |

**Table S7: Statistical variation in HPLC analysis.** To evaluate the intrinsic error of the HPLC (*Agilent*), a 10 ml solution of 55.5 mM sodium acetate was prepared in water and then injected into the HPLC six times. The consensus signal intensity is defined as exactly 555 µmol (55.5 mM). Evaluations of acetate concentration within 10 ml broth cultures were determined to be accurate to ± 2.2 µmol (0.22 mM).

| Replicate Number | Peak Area (unitless) | Acetate (mM) | Acetate (µmol) |
|---|---|---|---|
| 1 | 983073 | 55.69 | 556.9 |
| 2 | 983990 | 55.74 | 557.4 |
| 3 | 978185 | 55.41 | 554.1 |
| 4 | 981409 | 55.60 | 556.0 |
| 5 | 978268 | 55.42 | 554.2 |
| 6 | 973352 | 55.14 | 551.4 |
| **Average** | 979713 | 55.5 | 555 |
| **Std. Dev.** | 3933 | 0.22 | 2.2 |
| **Error (%)** | 0.40 | 0.40 | 0.40 |

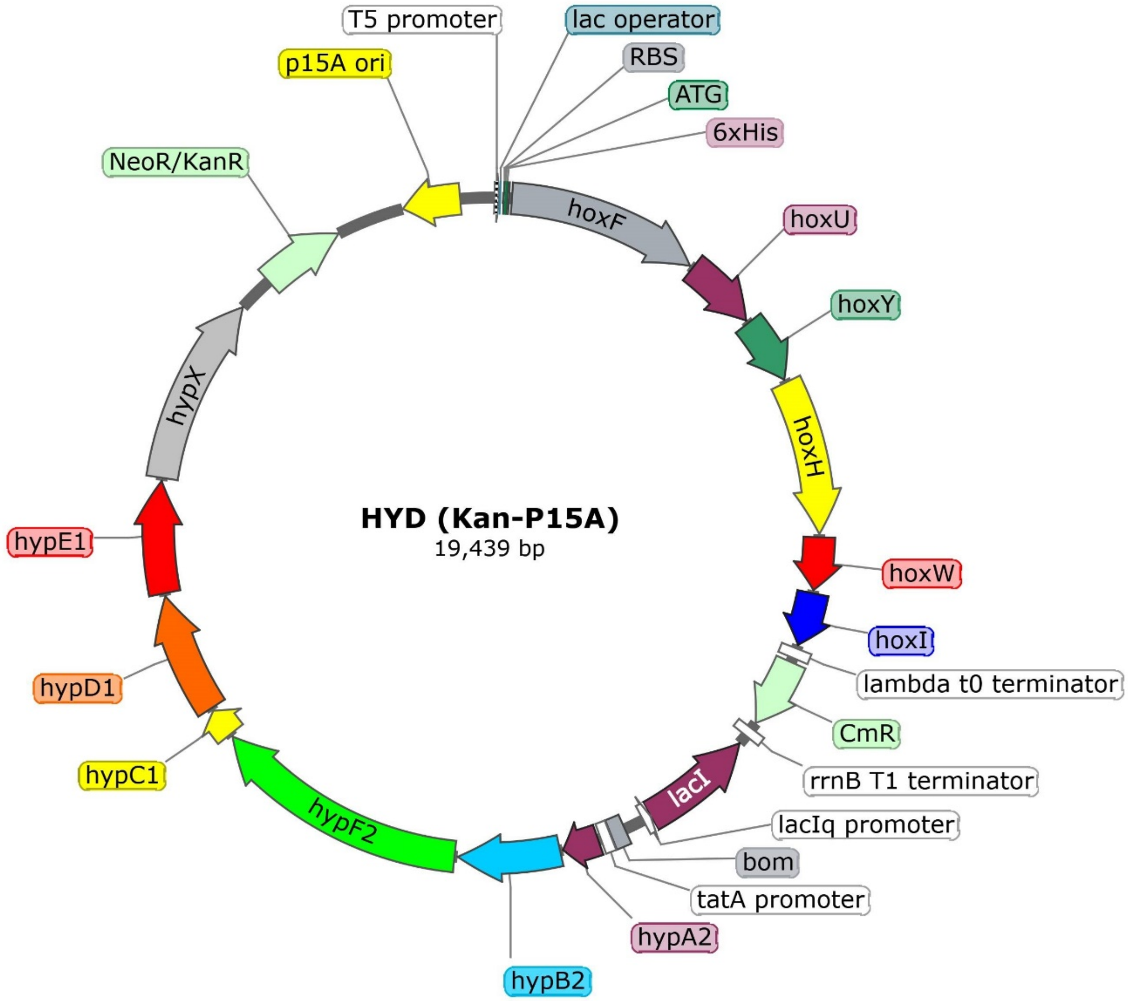


**Figure S5: Enlarged plasmid map of HYD (Kan-p15A).** The first operon encodes the hydrogenase heterotetramer (*hoxFUYH*) and maturation genes (*hoxWI*), which are inducible by IPTG via the T5 promoter. The second operon encodes the remaining maturation genes (*hypABCDEFX*) and is constitutively expressed via the *tatA* promoter. The chloramphenicol resistance marker ("CmR") is nonfunctional and is an artifact of the construction scheme (Figure S1 in *SI*).

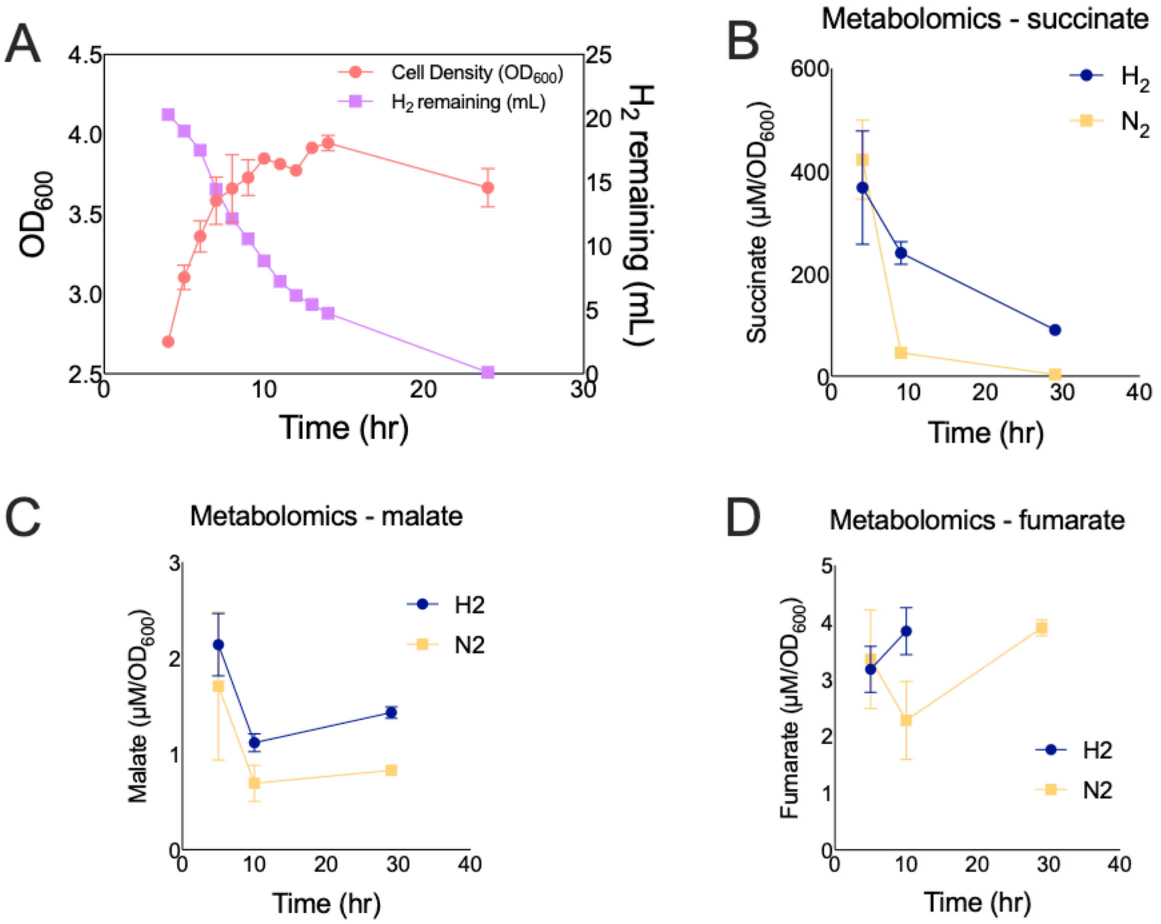


**Figure S6: Time-course experiment evaluating $H_2$ uptake in BW-HYD when grown in LB media and its effects on production of succinic acid, malate, and fumarate. (A)** Time-course experiment evaluating the growth profile and $H_2$ consumption of BW-HYD in LB media, using a simplified version of the scheme illustrated in Figure S3 (n = 2), for the purpose of designing a follow-up experiment assaying intracellular metabolite at key time-points. One gas reading at 11h and another at 14h were lost due to a handling error. No parallel experiment involving $N_2$-treatment was performed. **(B, C, D)** A follow-up experiment, designed based on the results of (A) and performed using fresh cultures on a different day, in which technical replicate cultures of BW-HYD were cultivated again in LB media, induced with IPTG at 4h and treated with 20 ml of $H_2$ in air (treatment) or 20 ml of $N_2$ in air (control). Intracellular concentrations of succinic acid, malate, and fumarate were evaluated at three time points: 4h (onset of $H_2$ treatment), 9h (active $H_2$ uptake), and 29h (early stationary phase). No data for fumarate at 29h ($H_2$-treated) was recorded. The optical density and hydrogen gas uptake at these three time points were also monitored by spectrophotometry and gas chromatography, respectively. The resulting data corresponded closely with the time-course results shown in panels A and B (data not shown). See *Prototyping the C. necator soluble hydrogenase (HYD) and Pseudomonas sp. 101 formate dehydrogenase (FDH)* in the *Main Text*.

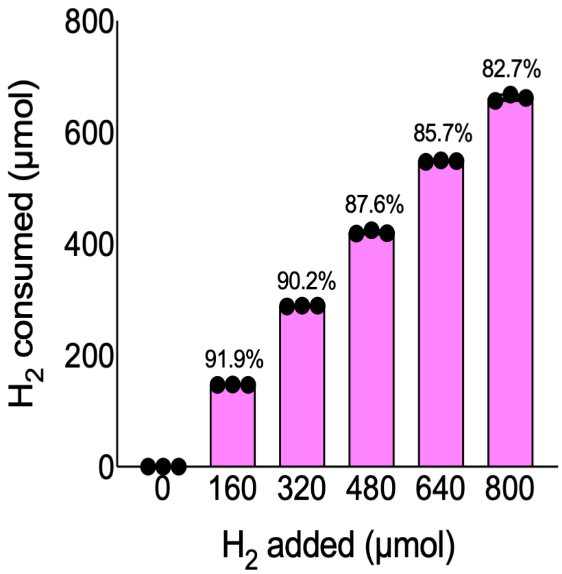


**Figure S7: $H_2$ uptake (µmol) in BW-HYD.** Refer to *Figure 2* and note that e*lectron donors prevent the decarboxylation of an organic carbon feedstock* in the Main Text. The X-axis represents the total hydrogen gas added, while the Y-axis shows the total hydrogen gas oxidized. Percentages indicate the fraction of added $H_2$ that was consumed following an overnight incubation. We speculate that the declining percentage of $H_2$ that is oxidized by BW-HYD associated with the rising initial volume of $H_2$ could be caused by saturation of the hydrogenase or by mass transfer limitations.

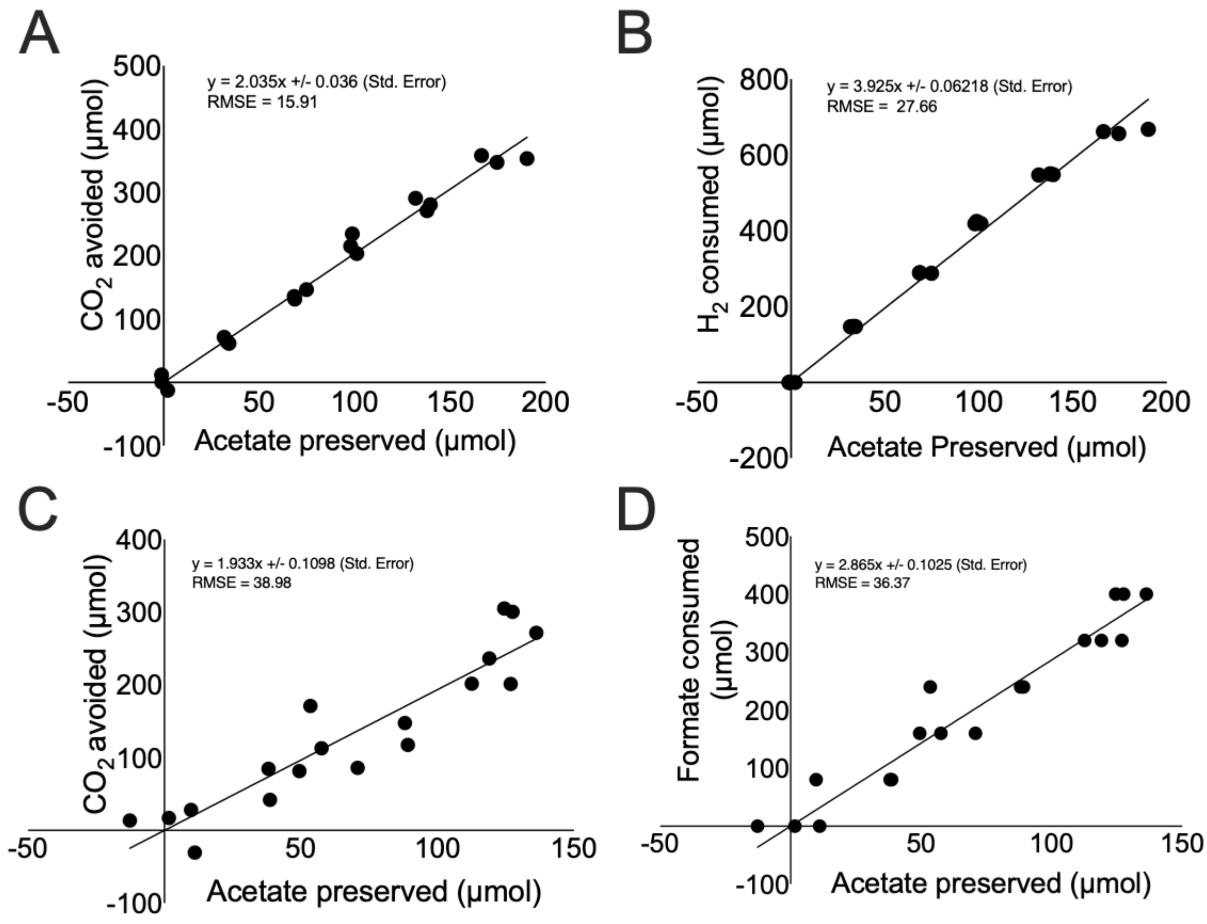


**Figure S8: Linear correlations observed between feedstock retention, CO2 evolution, and external energy carrier consumption. (A)** Moles of $CO_2$ abated by $H_2$ uptake versus moles of acetate retained by $H_2$ uptake (moles / moles), where a slope of two is expected based on stoichiometry. **(B)** Moles of $H_2$ consumed versus moles of acetate preserved (moles / moles), where the slope acts as a metric for the efficiency of hydrogen gas usage, with the lowest theoretical slope being 3.4 (see *Flux balance analysis to evaluate efficiency of hydrogen gas and formate usage* in the *Main Text*). **(C)** Moles of biogenic $CO_2$ avoided by formate uptake versus moles of acetate retained by formate uptake (moles / moles), with a slope of two expected according to stoichiometry. **(D)** Moles of formate consumed versus moles of acetate preserved (moles / moles), where the slope serves as a metric for the efficiency of formate usage, and the lowest theoretical slope is 2.82 (discussed in *Flux balance analysis to evaluate the efficiency of hydrogen gas and formate usage* in the *Main Text*). RMSE = Root Mean of the Standard Error.

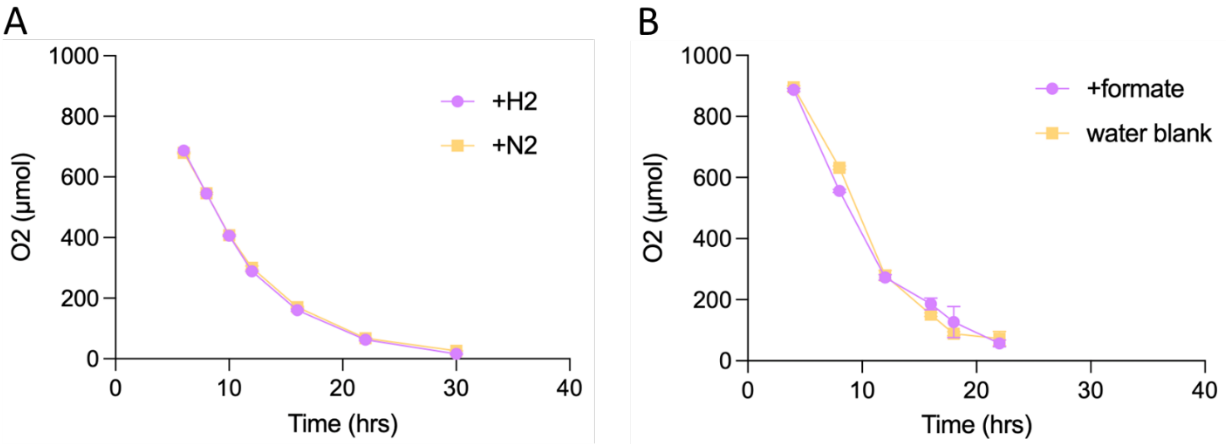


**Figure S9: Oxygen gas ($O_2$) consumption profile (µmol) by BW-HYD (A) and BW-FDH (B) when cultivated in EZ-Acetate.** See Figure 2C and Figure 3C in *main text*.

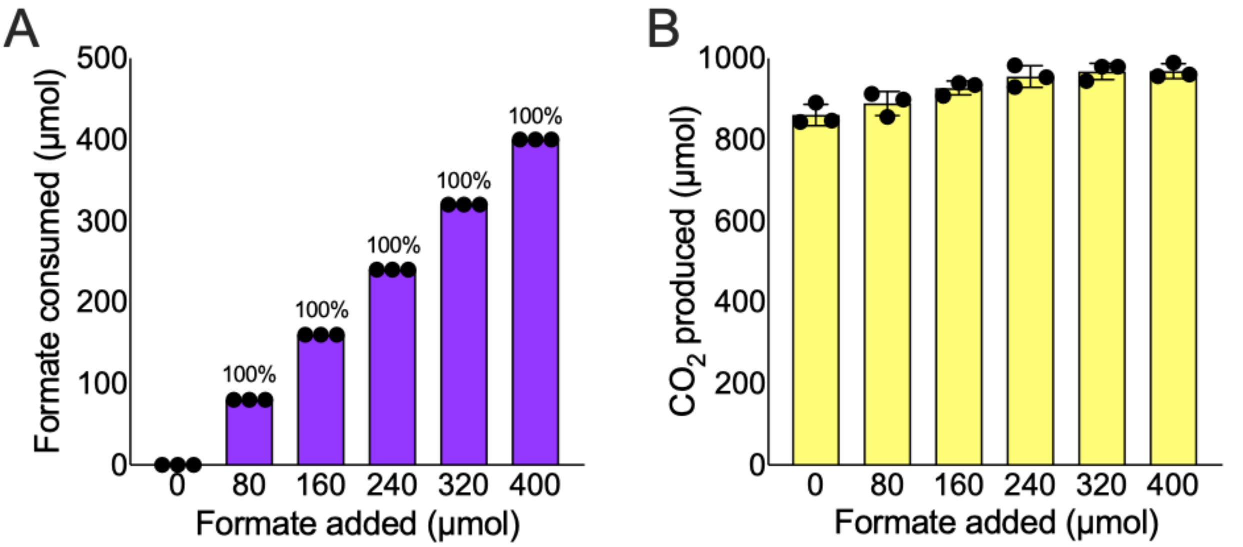


**Figure S10: Formate uptake (Left; µmol) and total $CO_2$ in headspace (Right; µmol) in BW-FDH.** Refer to *Figure 3* and the section "*Electron donors prevent the decarboxylation of an organic carbon feedstock"* in the *Main Text*. **(A)** The X-axis indicates total formate added, while the Y-axis displays total formate oxidized. The percentages represent the proportion of added formate consumed after an overnight incubation. We found that formate was undetectable in the media following this incubation, which we interpreted as total consumption of formate. **(B)** By stoichiometry, the oxidation of each mole of formate generates one mole of $CO_2$. Therefore, to calculate biogenic $CO_2$ in Figure 3 (*Main Text*), one must subtract the formate consumed from the total $CO_2$.

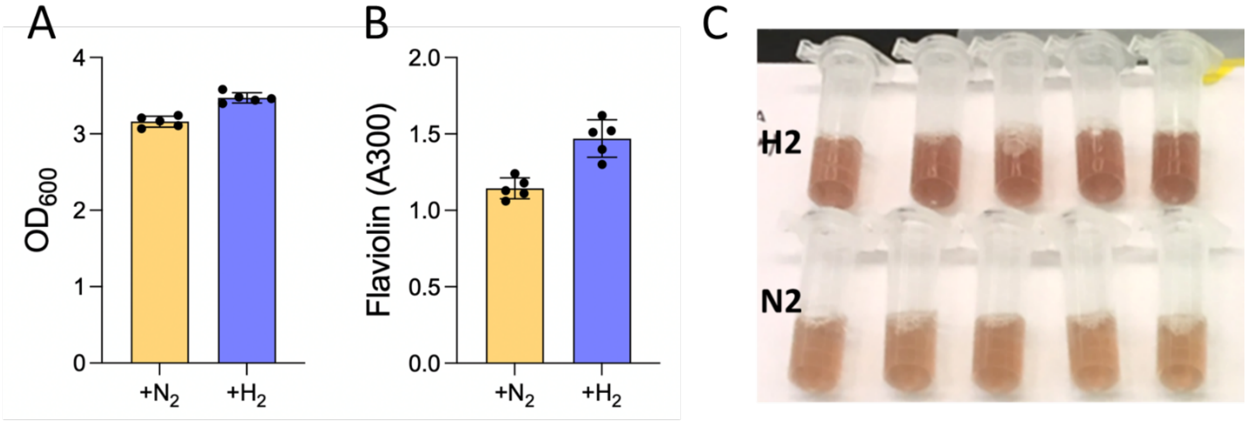


**Figure S11: Enhanced flaviolin production enabled by $H_2$ in BW-HYD-RppA**. Flaviolin is a non-reduced red pigment polyketide produced by the polyketide synthase RppA via the condensation of five units of malonyl-CoA. Oxidation by molecular oxygen of the unstable and colorless 2,4,6,8-tetrahydroxynaphthalene intermediate to flaviolin is spontaneous. **(A and B)** Final optical density (OD600) and relative flaviolin ($\lambda_{max}$ 300 nm) production observed in cultures of BW-HYD-RppA when grown overnight in LB media in the presence of 20% $H_2$ in air ("$H_2$") or 20% $N_2$ in air ("$N_2$"). **(C)** Photograph of broth samples.

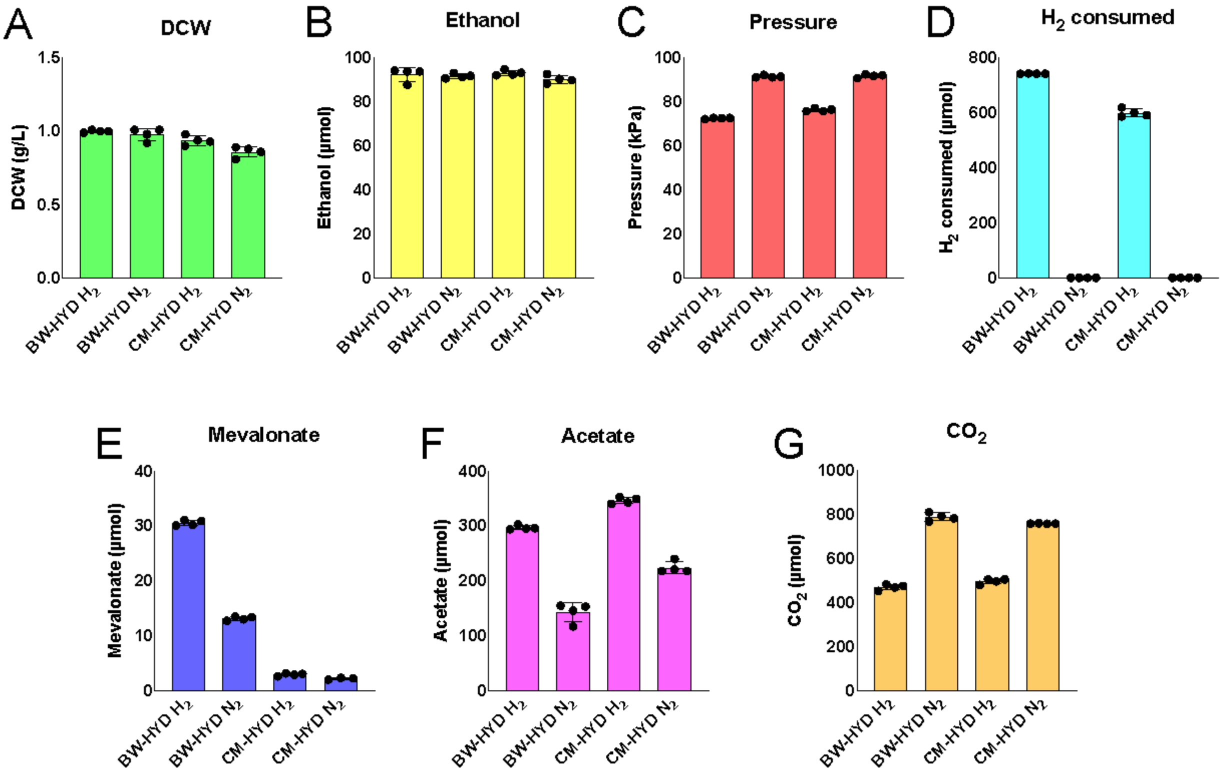


**Figure S12: Mevalonate titer achieved using EZ-Acetate in BW-HYD-MEV and CM-HYD-MEV**. We assessed mevalonate biosynthesis in BW25113 *E. coli* (BW-HYD-MEV) and CM15 *E. coli* (CM-HYD-MEV), both of which express *C. necator* hydrogenase, using the same scheme illustrated in Figure S2A. The only difference was that *E. coli* was transformed with HYD (Kan|P15A) and treated with $H_2$ or $N_2$ in air before being cultivated in EZ Rich – Acetate (55.5 mM). This experiment is interpreted as follows: Aside from a slight increase in biomass formation in CM15 *E. coli* treated with $H_2$ **(A)**, biomass formation and ethanol fermentation **(B)** are independent of $H_2$ uptake, thereby normalizing cellular energy needs. Internal bottle pressure indicates successful $H_2$ uptake **(C)**, which was directly confirmed when the internal gas was analyzed by gas chromatography **(D)**. Analysis of the mevalonate content by LC-MS revealed that $H_2$ uptake indeed increased mevalonate production, doubling the titer produced by BW-HYD from 13 mg/L to 30 mg/L, with a minimal 1 mg/L increase for CM-HYD **(E)**. However, the additional mevalonate produced from $H_2$ treatment (approximately 1.35 µmol) only requires cellular energy derived from the oxidation of about 20 µmol of $H_2$ (see *Electron donors provide exceptional cellular energy efficiency* in *Main Text* for explanation), suggesting that most of the 600 to 750 µmol of $H_2$ consumed by *E. coli* was directed toward other metabolic processes. Further investigation indicated that this energy helped prevent the decarboxylation of acetate **(F)**, as evidenced by the absence of $CO_2$ production **(G)**, in a manner that replicates the findings shown in *Electron donors prevent the decarboxylation of a carbon feedstock* in the *Main Text*. Producing mevalonate in *E. coli* via acetate feedstock is well-known to be metabolically bottlenecked due to limitations in acetyl-CoA supply (Jeung et al., 2023). Therefore, we surmised that this protocol was inadequate for demonstrating how external reducing power can enhance bioproduction, necessitating a glucose-based minimum medium.

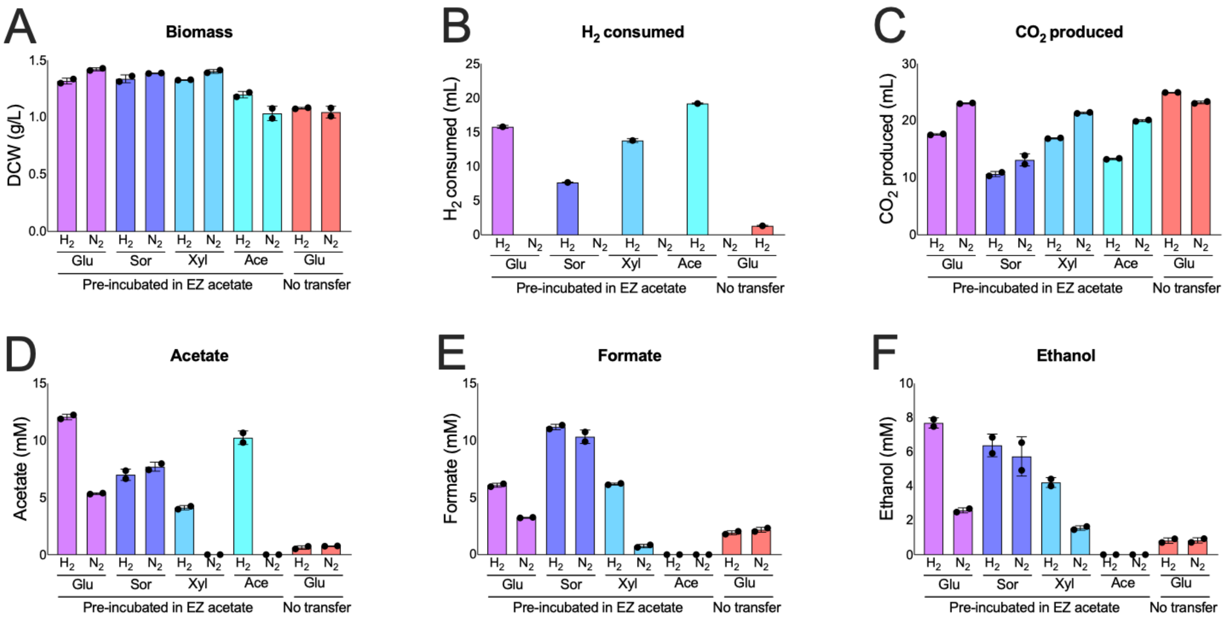


**Figure S13**: **Development of a pre-incubation protocol to enhance hydrogen gas uptake in BW-HYD when cultivated in various minimal media. (A)** Culture density (DCW; converted from OD600 readings using the consensus ratio); **(B)** Hydrogen gas consumed by *E. coli* (ml), with an initial provision of 20 ml of $H_2$; **(C)** $CO_2$ production (ml); **(D, E, F)** Acetate, formate, and ethanol detected in the culture supernatant (mM). Samples labeled "Pre-incubated in EZ acetate" were treated similarly to the scheme illustrated in Figure S4A, except that the pre-cultivation media was EZ Rich with 55.5 mM acetate as the primary carbon source, and the transfer media was M9 minimal media containing either 27.75 mM glucose, 27.75 mM sorbitol, 27.75 mM xylose, or 55.5 mM acetate as the sole carbon source. Samples labeled "No Transfer" were processed in a manner similar to the scheme illustrated in Figure S4B, with the exception that *E. coli* was expressing *C. necator* hydrogenase, and cells were provided with 20 ml of $H_2$ in air or 20 ml of $N_2$ in air. No glucose, sorbitol, or xylose was detected in the culture supernatant, and the presence of glycerol, lactate, and succinate was negligible (data not shown). See *Electron donors increase mevalonate yield with optimizable efficiency* in the *Main Text*.

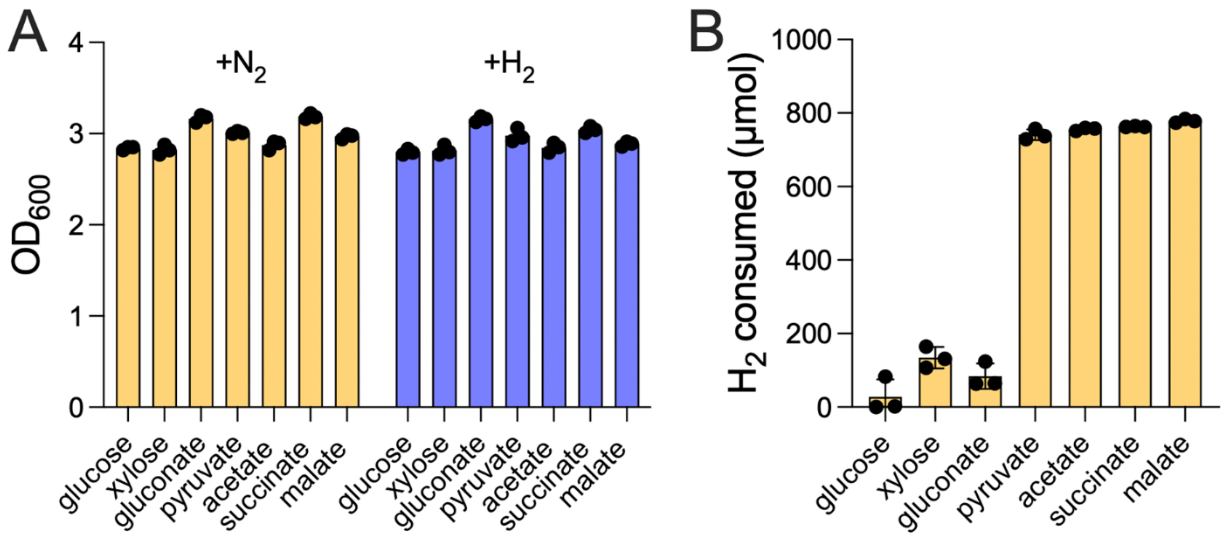


**Figure S14: Hydrogen gas uptake by BW-HYD in sugars and oxidized feedstocks. (A)** Final culture density (OD600) and **(B)** $H_2$ consumption (μmol) following an overnight procedure (similar to Figure S2A) wherein BW-HYD was cultivated in EZ Rich containing either 27.75 mM glucose, 27.75 mM xylose, 27.75 mM gluconate, 55.5 mM pyruvate, 55.5 mM acetate, 27.75 mM succinate, or 27.75 mM malate, in stoppered serum vials with a headspace initially composed of 800 μmol $H_2$ in air (+) or an equivalent volume of $N_2$ in air (-). Performed in triplicate (n=3).

**Table S8: Micromoles (µmol) of mixed acid products (formate, ethanol, acetate, glycerol, lactate, succinate) observed in the culture supernatant (10 ml) of BW-FDH-MEV, CM-FDH-MEV, BW-HYD-MEV, and CM-HYD-MEV.** Refer to Figures 4D and 4I in the *Main Text*. *Compared* to *E. coli* BW25113, the observed decreases in acetate and increases in formate in hydrogenase-laden strains, along with the decrease in acetate in formate dehydrogenase-laden strains, correspond to the metabolic profile of *ΔackAΔpta E. coli*. The presence of ethanol in CM15 is intriguing because this strain contains a *ΔadhE* knockout. We speculate that the concurrent knockout of multiple primary fermentation pathways forces ethanol fermentation through secondary alcohol dehydrogenases, as noted elsewhere (Yang et al., 1999; Zhu et al., 2011). ND = Not detected

| | BW-FDH-MEV | | CM-FDH-MEV | | BW-HYD-MEV | | CM-HYD-MEV | |
|---|---|---|---|---|---|---|---|---|
| | (-) | (+) | (-) | (+) | (-) | (+) | (-) | (+) |
| Formate | ND | ND | ND | ND | 7.2 ±0.5 | 12.2 ±1.6 | 74.5 ±2.3 | 82.7 ±3.4 |
| Acetate | 24.4 ±1.0 | 42.9 ±0.8 | 0.28 ±0.01 | 12.3 ±1.0 | 72.3 ±4.0 | 109.9 ±4.3 | 26.6 ±0.4 | 26.5 ±0.1 |
| Ethanol | 171.7 ±1.7 | 167.8 ±5.4 | 165.6 ±2.5 | 171.8 ±5.6 | 119.7 ±4.4 | 139.2 ±5.5 | 100.6 ±0.1 | 101.5 ±0.4 |
| Glycerol | 3.6 ±0.3 | 4.0 ±0.3 | 6.2 ±0.4 | 4.81 ±0.03 | ND | ND | ND | ND |
| Lactate | ND | ND | ND | ND | ND | ND | ND | ND |
| Succinate | 0.82 ±0.04 | 0.89 ±0.02 | 1.17 ±0.05 | 1.24 ±0.01 | ND | ND | ND | ND |

**Supplemental References**

*Note*: Works cited in *Supporting Information* not found below are found in the *Main Text*.


1. Ahrné, E., Molzahn, L., Glatter, T., Schmidt, A. Critical assessment of proteome-wide label-free absolute abundance estimation strategies. Proteomics 13, 2567-2578 (2013).
2. Amer, B., Kakumanu, R., Baidoo, E.E.K. HILIC-MS analysis of central carbon metabolites in gram-negative bacteria. Protocols.io https://dx.doi.org/10.17504/protocols.io.4r3l2opzxv1y/v1
3. Chen, Y., Gin, J.W., Wang, Y., de Raad, M., Tan, S., Hillson, N.J., Northen, T.R., Adams, P.D., Petzold, C.J. Alkaline-SDS cell lysis of microbes with acetone protein precipitation for proteomic sample preparation in 96-well plate formate. PLoS ONE 18, e0288102 (2023).
4. Deng, Y., Beahm, D.R., Ionov, S., Sarpeshkar, R. Measuring and modeling energy and power consumption in living microbial cells with a synthetic ATP reporter. BMC Biol. 19, 101 (2021).
5. Ebrahim, A., Lerman, J.A., Palsson, B.O., Hyduke, D.R. COBRApy: COnstraints-Based Reconstruction and Analysis for Python. BMC Syst. Biol. 7, 74 (2013).
6. King, Z.A., Lu, J., Dräger, A., Miller, P., Federowicz, S., Lerman, J.A., Ebrahim, A., Palsson, B.O., Lewis, N.E. BiGG models: A platform for integrating, standardizing and sharing genome-scale models. Nucleic Acids Res. 44, D515-D522 (2016).
7. Klamt, S., Müller, S., Regensburger, G., Zanghellini, J. A mathematical framework for yield (vs. rate) optimization in constraint-based modeling and applications in metabolic engineering. Metab. Eng. 47, 153-169 (2018).
8. Kumari, S., Beatty, C.M., Browning, D.F., Busby, S.J., Simel, E.J., Hovel-Miner, G., Wolfe, A.J. Regulation of acetyl coenzyme A synthetase in *Escherichia coli*. J. Bacteriol. 182, 4173–4179 (2000).
9. Lauterbach, L., Lenz, O. Catalytic production of hydrogen peroxide and water by oxygen-tolerant [NiFe]-hydrogenase during H2 cycling in the presence of O2. J. Am. Chem. Soc. 135, 17897–17905 (2013).
10. Milo, R., Phillips, R. *Cell Biology by the Numbers.* CRC Press (2015).
11. Orth, J.D., Fleming, R.M.T., Palsson, B.Ø. Reconstruction and use of microbial metabolic networks: The core *Escherichia coli* metabolic model as an educational guide. EcoSal Plus 4 (2010).
12. Perez-Riverol, Y., Bai, J., Bandla, C., García-Seisdedos, D., Hewapathirana, S., Kamatchinathan, S., Kundu, D.J., Prakash, A., Frericks-Zipper, A., Eisenacher, M., Walzer, M., Wang, S., Brazma, A., Vizcaíno, J.A. The PRIDE database resources in 2022: A hub for mass spectrometry-based proteomics evidences. Nucleic Acids Res. 50, D543-D552 (2021).
13. Sauer, U., Canonaco, F., Heri, S., Perrenoud, A., Fischer, E. The soluble and membrane-bound transhydrogenases UdhA and PntAB have divergent functions in NADPH metabolism of *Escherichia coli*. J. Biol. Chem. 279, 6613-6619 (2004).
14. Silva, J.C., Gorenstein, M.V., Li, G.Z., Vissers, J.P.C., Geromanos, S.J. Absolute quantification of proteins by LCMSE: A virtue of parallel MS acquisition. Mol. Cell. Proteom. 5, 144-56 (2006).
15. Wang, G., Kakumanu, R., Amer, B., Turumtay, E.A., Baidoo, E.E.K. Reversed phase LC-MS analysis of organic acids involved in the tricarboxylic acid cycle V.2. *protocols.io* (2025) dx.doi.org/10.17504/protocols.io.bp2l6dz15vqe/v2

16. Wenk, S., Schann, K., He, H., Rainaldi, V., Kim, S., Lindner, S.N., Bar-Even, A. An "energy-auxotroph" *Escherichia coli* provides an in vivo platform for assessing NADH regeneration systems. Biotechnol. Bioeng. 117, 3422-3434 (2020).
17. Yang, Y. T., San, K. Y. & Bennett, G. N. Redistribution of metabolic fluxes in Escherichia coli with fermentative lactate dehydrogenase overexpression and deletion. *Metab. Eng.* **1**, 141–152 (1999).
18. Zhu, H., Gonzalez, R. & Bobik, T. A. Coproduction of acetaldehyde and hydrogen during glucose fermentation by Escherichia coli. *Appl. Environ. Microbiol.* **77**, 6441–6450 (2011).
